# The 3D genome of pediatric B-cell precursor acute lymphoblastic leukemia

**DOI:** 10.1101/2025.07.22.666073

**Authors:** Larissa H Moura-Castro, Efe Aydın, Gladys Telliam Dushime, Eleanor L Woodward, Anders Castor, Rebeqa Gunnarsson, Henrik Lilljebjörn, Linda Olsson-Arvidsson, Thoas Fioretos, Bertil Johansson, Marcus Järås, Anna K Hagström-Andersson, Minjun Yang, Kajsa Paulsson

## Abstract

Whereas the molecular pathogenesis of childhood B-cell precursor acute lymphoblastic leukemia (BCP ALL) has been studied extensively, its 3D chromatin landscape – of vast importance for gene regulation – remains poorly explored. Here, we applied Micro-C, a high-resolution variant of Hi-C, to 35 primary pediatric BCP ALL cases, spanning all major genetic subtypes. We present a complete view of the chromatin interaction landscape in childhood ALL, with resolutions reaching up to 5 kb in individual samples and 1 kb in the aggregated dataset. Somatic genetic aberrations – including fusion genes, aneuploidy, and structural variants – were found to profoundly reshape the 3D genome organization, impacting chromatin compartmentalization (A/B), topologically associating domain (TAD) architecture, and regulatory element positioning. Notably, chromosomal gains were associated with weakened TAD boundaries and widespread gene dysregulation. In addition, our analysis identified over 25,000 chromatin loops anchored at regulatory elements—e.g., enhancer–promoter loops—regulating the expression of more than 10,000 protein-coding genes. Among these, we highlight regulatory loops that drive gene expression differences between BCP ALL subtypes in the absence of concurrent somatic genetic aberrations, including the known driver genes *HOXA9, FLT3, TP53, CD44, IKZF1, ERG,* and *XBP1*. Taken together, our study gives unprecedented insights into chromatin organization and gene regulation in the leukemogenesis of BCP ALL.

## Introduction

B-cell precursor acute lymphoblastic leukemia (BCP ALL) is characterized by abnormal proliferation of B-lymphoid progenitor cells and is the most common malignancy in children^1^. It is divided into clinically important genetic subtypes, with high hyperdiploid (HeH; 51-67 chromosomes) and *ETV6*::*RUNX1* cases comprising approximately half of the cases. Other major subtypes include cases with *TCF3*::*PBX1* or *BCR*::*ABL1* fusion genes, rearrangements of *KMT2A* (*KMT2A*-r; previously *MLL*), or *DUX4* (*DUX4*-r), intrachromosomal amplification of chromosome 21 (iAMP21), and cases characterized by specific chromosome numbers such as near-haploidy (24-30 chromosomes)^1^. The overall prognosis of pediatric BCP ALL is favorable on current treatment protocols, but certain subgroups still fare poorly. Furthermore, studies to deescalate treatment and minimize toxic side effects are ongoing. Overall, an improved biological understanding is needed for this malignancy^1^.

Whilst the molecular pathogenesis of various BCP ALL genetic subtypes has been studied extensively, their 3D genome landscape remains poorly explored. Chromatin folding is a hierarchical process where chromosomes first occupy distinct territories in the nucleus, followed by organization of compartments that cluster predominately active (compartment A) or inactive (compartment B) genes^2^. Topologically associating domains (TADs) up to 1 Mb in size are then formed, with TAD boundaries being physical barriers characterized by binding of CTCF and cohesin promoting intra-interactions while insulating the region from neighboring chromatin^2,3^. High-throughput chromosome conformation capture (Hi-C) has enabled the study of not only A/B compartments and TADs but also finer, intra-TAD structures, including chromatin loops between regulatory elements such as enhancers and promoters on a genome-wide scale^2^. The 3D chromatin architecture plays a critical role in gene regulation across all these levels, making its exploration crucial for our understanding of transcription control and, by extension, tumorigenesis.

Here, we utilize Micro-C – a variant of Hi-C with nucleosome-level resolution^2^ – whole genome sequencing (WGS), and RNA-sequencing to explore the 3D genome landscape of childhood BCP ALL primary patient samples, including all major genetic subtypes. We show that somatic genetic events lead to specific chromatin states, including position effects from structural rearrangements, discover weak TAD boundaries associated with gene dysregulation in aneuploid cases, and present very high-resolution data of enhancer-promoter interactions affecting the expression of more than 10,000 protein coding genes in BCP ALL, including well-known leukemic and oncogenic drivers such as *IKZF1*, *FLT3*, and *TP53*, providing novel insights into BCP ALL leukemogenesis.

## Results

### Somatic aberrations profoundly impact chromatin organization in pediatric BCP ALL

To investigate 3D chromatin conformation changes among different genetic subtypes of BCP ALL, we performed Micro-C on 35 BCP ALLs and one mixed phenotype acute leukemia (MPAL), resulting in an average of 1.6 billion unique valid sequence tags per sample (**Supplementary tables 1 and 2**). Chromatin interactions were analyzed at different organizational levels, starting with overall compartmentalization of the genome into predominantly active (A compartment) or inactive (B compartment) regions and TAD structure^2^. First, we compared primary BCP ALL with normal hematopoietic cells, comprising Micro-C analysis on sorted pro-/pre-B cells obtained from CD34+ enriched mononuclear cord blood and one ALL remission sample in addition to publicly available Hi-C data from four peripheral blood mononuclear cell (PBMC) samples and three CD34+ hematopoietic stem cell (HSPC) samples^4^ (**Supplementary table 2**). Principal component analysis (PCA) of A/B compartment 500 kb bins revealed three separate clusters: one consisting of the HSPC, the pro-B/pre-B and *KMT2A*-r samples, one comprising the PBMC and remission samples, and one more diffuse cluster containing the remaining primary leukemia samples (**Supplementary figure 1**). Comparing the number of A compartment bins, which corresponds to the amount of open, more active chromatin, we found that primary patient samples and HSPC/pro-/pre-B cells had a significantly smaller part of the genome in the A compartment (median 2,454 and 2,509, respectively) than the more mature PBMC/remission cells (median 2,878; two-sided Mann-Whitney U test; *P*=0.001 and *P*=0.016, respectively) (**Figure 1A**). Thus, primary BCP ALL is more similar to immature hematopoietic cells and there are genome-wide differences between genetic subtypes in their overall chromatin organization.

**Figure 1.**
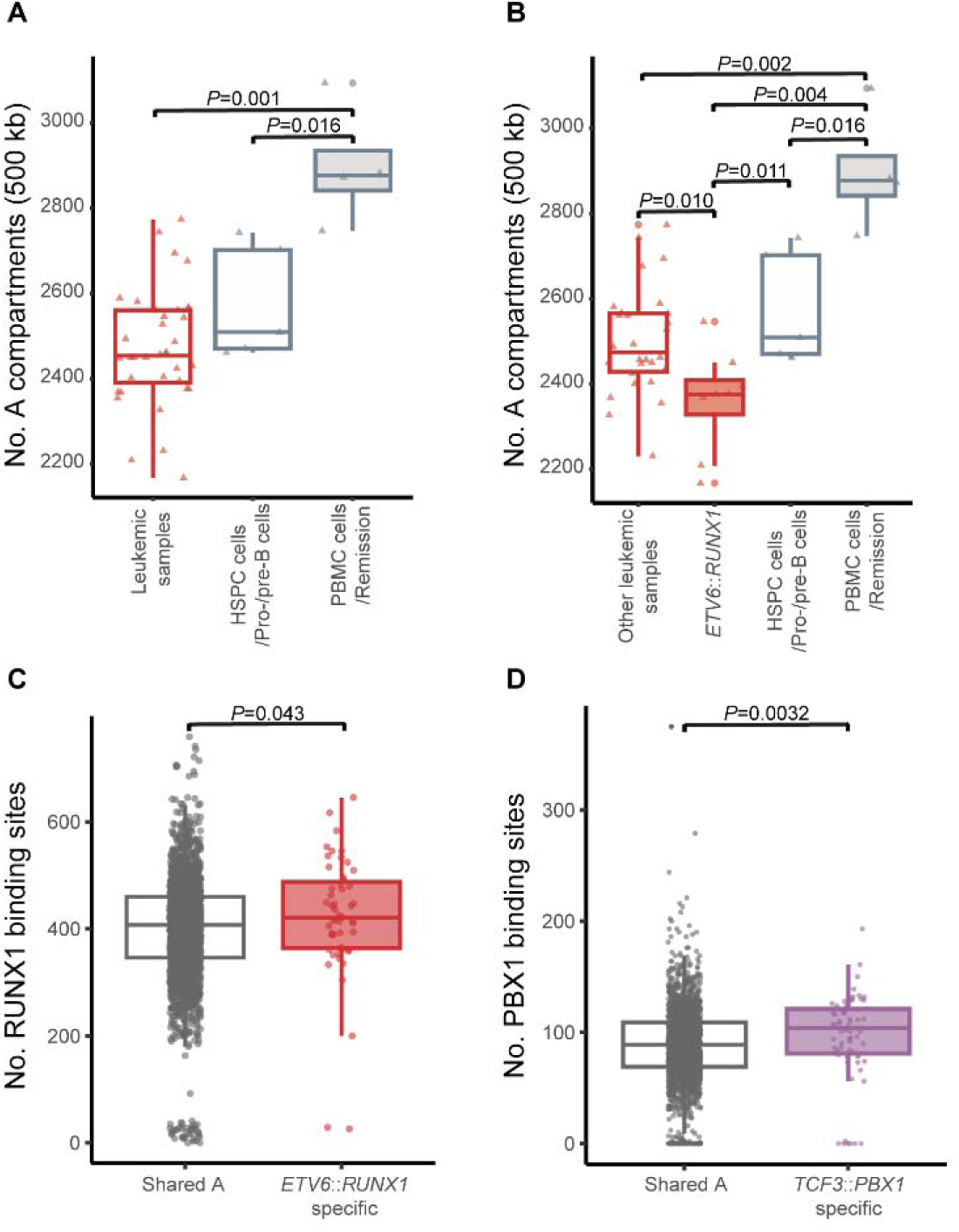
Analysis of A/B compartments in B-cell precursor acute lymphoblastic leukemia (BCP ALL). Open chromatin profiles of leukemic subtypes and normal cells measured as total number of 500 kb bins in open chromatin state (A compartment). **A.** Amount of open chromatin in normal cells is higher than in leukemic cells (two-sided Mann-Whitney test). **B.** *ETV6*::*RUNX1* cases have significantly less open chromatin compared to the other BCP ALL cases combined and to the normal cells (two-sided Mann-Whitney test). **C-D.** Number of predicted transcription factor binding sites on A compartments, separated by those shared across all subgroups and those specific to the indicated fusion, showing enrichment of sites for the respective transcription factor. (C) RUNX1 in *ETV6::RUNX1*; (D) PBX1 in *TCF3::PBX1.* Abbreviations: HSPC, hematopoietic stem and precursor cells; PBMC, peripheral blood mononuclear cells.

Turning to genetic subtypes, *ETV6*::*RUNX1* cases had significantly less open chromatin compared with all the other BCP ALL cases combined (median 2,376 vs 2,474; two-sided Mann-Whitney U test; *P*=0.01) and with both the HSPC and PBMC clusters (two-sided Mann-Whitney U test; *P*=0.011 and *P*=0.004, respectively) (**Figure 1B**). Integrating with RNA-seq data, we could see that A-to-B shifts between subtypes in general were associated with decreased gene expression and B-to-A shifts with increased gene expression (**Supplementary figure 2, Supplementary tables 3 and 4**), in line with the A compartment corresponding to more open chromatin and with previous studies showing that B-to-A shifts are associated with increased gene expression in different cell types^5–7^. To see whether shifts to the A compartment were associated with activity of transcription factors specific for genetic subtypes, we analyzed A compartment sequences specific to *ETV6*::*RUNX1* and *TCF3*::*PBX1* for enrichment of RUNX1 and PBX1 binding motifs, respectively. Both subtypes exhibited a significantly higher number of predicted binding sites for their corresponding transcription factors compared to shared A compartments (**Figure 1C, D, Supplementary table 3**), underscoring the interplay between somatic genetic events and chromatin organization.

Next, we investigated the TAD organization in pediatric BCP ALL. Enhancer-promoter interactions are diminished across TAD boundaries, giving TAD organization a pivotal role in overall gene regulation^8^. At 25 kb resolution, we identified 12,635 TAD boundaries with a median TAD size of approximately 300 kb (**Supplementary tables 2 and 5**). PCA of the boundary scores of common TAD boundaries present in at least half of the BCP ALL cases showed three clear clusters (**Figure 2A**). Cluster 1, the “aneuploid” cluster, comprised all cases with ≥50 chromosomes, including all HeH, the near-haploid case (that harbored a major duplicated clone with 54 chromosomes), a *BCR*::*ABL1* case with 51 chromosomes, and a B-other case with 50 chromosomes. *ETV6*::*RUNX1* cases and one *DUX4-r* case with a somatic *ETV6* deletion formed a second cluster (cluster 2). Cluster 3 comprised the four *TCF3*::*PBX1* cases, while the remaining cases were not part of any cluster. Comparative analysis of TAD boundary features revealed marked differences. Cluster 2 displayed the highest number of TAD boundaries (mean 8,310; range 7,843-8,618; *P*=0.00013, two-sided Mann-Whitney U test) compared to all other BCP ALL cases, followed by cluster 3 (mean 7,419; range 5,995-8,098). In contrast, cluster 1 exhibited a statistically significant reduction in boundary numbers (mean 7,380; range 5,814-8,207; *P*=5.6×10^-5^, two-sided Mann-Whitney U test) and had an increase in average TAD length by 30 kb (**Figure 2B, Supplementary table 2**). In line with this, analysis of TAD boundary strength showed that cluster 2 had the strongest boundary strength compared to the remaining BCP ALL cases (*P*=0.0001947, two-sided Mann-Whitney U test), indicating a more robust chromatin organization, whereas cluster 1 displayed a significant loss in boundary strength (*P*=0.003824, two-sided Mann-Whitney U test) (**Figure 2C, Supplementary table 5**).

**Figure 2.**
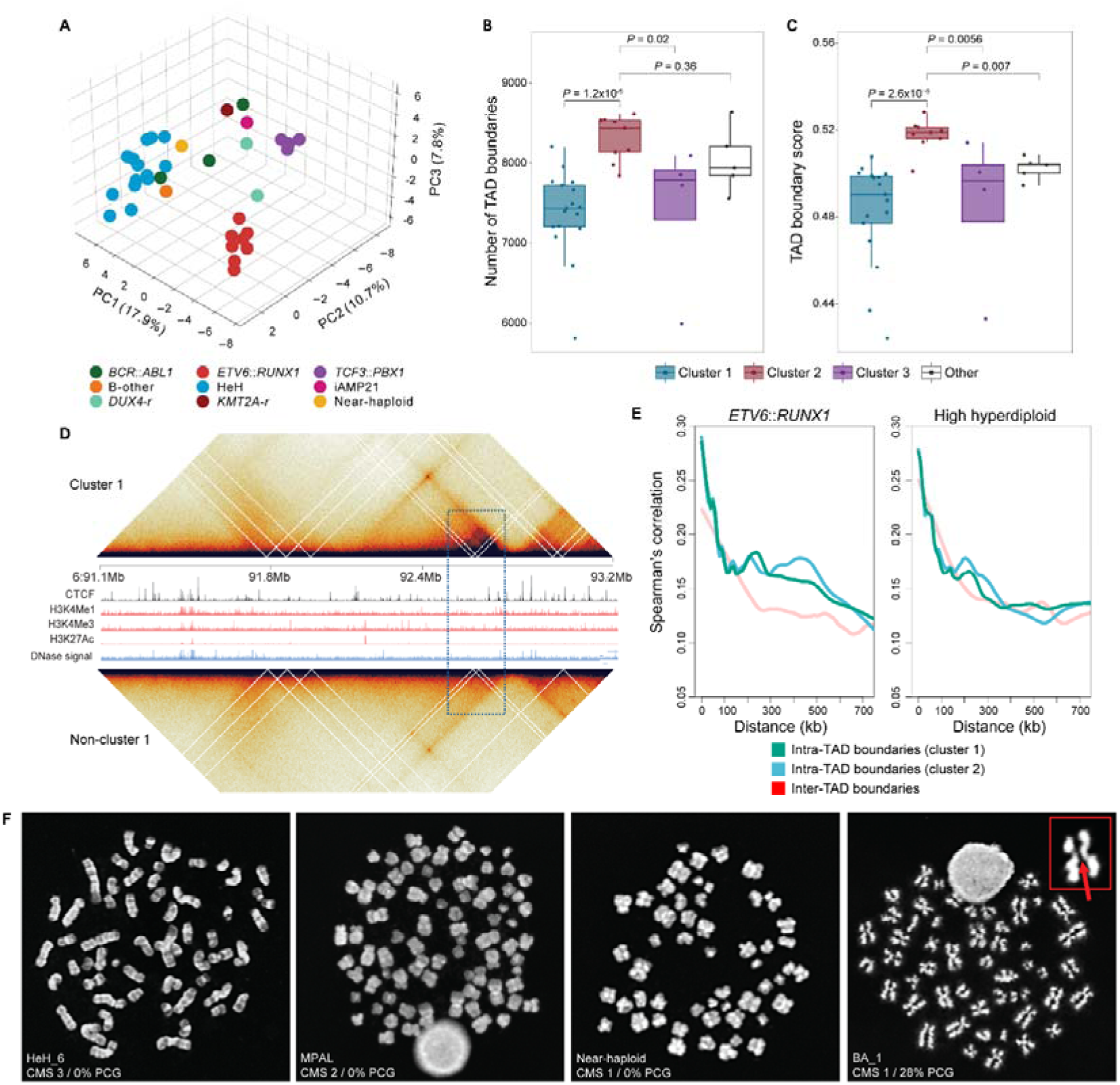
Topologically associating domain (TAD) analysis of 35 B-cell precursor acute lymphoblastic leukemia samples (BCP ALL). **A.** Principal component analysis (PCA) of topologically associating domain (TAD) boundary scores from 35 BCP ALL samples revealed three distinct clusters. Cluster 1 comprised all aneuploid BCP ALL cases, cluster 2 included all *ETV6*::*RUNX1* cases and one *DUX4*-rearranged case with a somatic *ETV6* deletion, and cluster 3 consisted of all *TCF3*::*PBX1* cases. **B.** Boxplots showing the number of TAD boundaries at 25 kb resolution. **C.** Boxplots of TAD boundary strength calculated using Chromosight at 25 kb resolution. **D.** ICE-normalized pooled Micro-C contact heatmaps at 25 kb resolution. The middle panels illustrate ChIP-seq tracks from the GM12878 cell line. A boundary gain event on chromosome 6 (blue square) was observed in cluster 1 cases compared to non-cluster 1 cases; this gained boundary lacked detectable CTCF or epigenetic marker binding signals based on the GM12878 ChIP-seq data. **E.** Spearman’s correlation analysis of the expression of gene pairs as a function of genomic distance revealed that in *ETV6*::*RUNX1* cases (left; n=47), gene pairs within TAD boundaries defined by cluster 2 (blue) demonstrated higher correlation than those within cluster 1 boundaries (green) or in different domains (red). In contrast, no significant differences were observed among high hyperdiploid samples (right; n=55), suggesting that the transcriptional dysregulation in cluster 1 cases is associated with TAD boundary organization. **F.** Examples of metaphase spreads classified according to chromosome morphology score (CMS) and percentage of cohesion defects (PCG), ranging from CMS 3 or good morphology and no PCG (far left) to CMS 1 or poor morphology and 28% PCG (far right) – where the zoomed-in chromosome (in red) shows in more detail the gap between sister chromatids.

Taken together, our analysis revealed that somatic genetic events profoundly impact chromatin organization. Notably, *ETV6::RUNX1* is associated with highly organized chromatin, with a larger proportion of the genome in the inactive B compartment and a higher number of TADs, whereas aneuploid cases, including the HeH subtype, show fewer and weaker TAD boundaries.

### Chromosomal gains lead to genome-wide changes in TAD structure in BCP ALL

The finding that all cases with ≥50 chromosomes displayed a similar TAD organization, regardless of the primary genetic event, is surprising, given that these represent different genetic subtypes with widely varying disease etiology and clinical features. To understand this phenomenon, we analyzed trisomic and disomic chromosomes separately, to see whether there was a direct effect on the TAD boundaries in gained chromosomes. However, no difference in boundary strength was observed, suggesting that the chromatin disorganization in these aneuploid cases is a genome-wide phenomenon and not associated with the copy number of a particular chromosome (**Supplementary figure 3A**).

CTCF binding is a main determinant of TAD boundary formation. Notably, we have previously shown that HeH ALL exhibit reduced mRNA and protein levels of *CTCF* ^9^. In line with this, cluster 1 cases in this study showed significantly lower *CTCF* expression compared to clusters 2 and 3, potentially contributing to the loss of TAD boundaries. Furthermore, regions exhibiting TAD boundary weakening in cluster 1 were significantly enriched for high-density CTCF binding sites, showing an odds ratio (high vs. low CTCF peak density) of 1.8, compared to 1.07 and 1.25 in clusters 2 and 3, respectively. This suggests disruption in CTCF-mediated insulation as a potential mechanism underlying TAD boundary weakening in cluster 1. Conversely, boundaries that gained strength in cluster 1 predominantly featured lower CTCF binding site density, with an odds ratio of 0.71 compared to 1.7 and 2.3 in clusters 2 and 3, respectively, indicating that strengthened boundaries in cluster 1 may emerge via mechanisms independent of strong CTCF binding (**Figure 2D, Supplementary figure 3B, 4**).

To understand how weaker TAD boundaries in aneuploid BCP ALL affect gene expression, we derived TAD boundaries from merged chromatin contact maps specifically from clusters 1 and 2 cases. A similar analysis with canonical TAD boundaries has previously shown genome-wide transcriptional dysregulation in HeH^9^. Briefly, we utilized RNA-seq data from 55 HeH and 47 *ETV6*::*RUNX1* cases and classified neighboring gene pairs based on their position relative to TAD boundaries as either inter-TAD (separated by a boundary) or intra-TAD (within the same TAD). Given the differences in TAD boundary structure between clusters 1 (including HeH) and 2 (including *ETV6::RUNX1*), we hypothesized that the expression of intra-TAD gene pairs should be more correlated when defined by the corresponding TAD data. Consistent with this, higher correlation scores for intra-TAD gene pairs were seen in *ETV6*::*RUNX1* when using cluster 2-derived TAD boundaries rather than those derived from cluster 1. In contrast, correlation scores for intra-versus inter-TAD gene pairs in HeH showed no difference regardless of whether TAD boundaries were derived from clusters 1 or 2 (**Figure 2E**). Taken together, this implies that the loss of TAD boundary integrity and diminished boundary strength seen in aneuploid ALL may lead to aberrant enhancer-promoter interactions across boundaries, thereby contributing to the dysregulation of gene expression seen in aneuploid cases and potentially causing leukemogenesis.

HeH BCP ALL cases have a high frequency of sister chromatid cohesion defects as well as poor chromosome morphology^9,10^. Considering our present findings that they also have fewer TADs and weaker TAD boundaries, we decided to investigate whether these different phenomena were associated in general in pediatric BCP ALL. Sister chromatid cohesion defects were analyzed by cytogenetic techniques and determined based on the presence of primary constriction gaps (PCGs) between the sister chromatids at metaphase (**Figure 2F**). The percentage of cells with cohesion defects ranged from 0 to 85% in the 35 cases analyzed (**Supplementary table 6**). In agreement with previous studies^10,11^, the HeH subtype displayed significantly higher levels of cohesion defects compared to non-HeH cases (median 14% vs. 4%, two-sided Mann-Whitney U test; *P*=0.0068) (**Supplementary figure 5A**). Chromosome morphology scoring (CMS) was also performed on cytogenetic preparations based on the appearance, condensation, and resolution of the metaphase chromosomes (**Supplementary table 6, Figure 2F**), using a scale of 1 to 3, with 1 corresponding to the poorest and 3 to the best chromosome resolution and morphology^9^. Overall, *ETV6*::*RUNX1* and *DUX4-*r cases displayed consistently good morphology (median CMS 2.14 and 2.49, respectively), whereas *TCF3*::*PBX1* cases and HeH cases displayed poorer chromosome morphology (median CMS 1.97 and 1.93, respectively) (**Supplementary figure 5B**). Both PCG percentage and CMS correlated with the number of TADs (*P*=0.0029 and *P*=0.018, respectively, Spearman’s correlation) (**Supplementary figure 5C**) but not with TAD boundary strength, with increased TAD numbers being associated with fewer cohesion defects and better chromosome morphology. However, when analyzing HeH and non-HeH cases separately, the correlations were statistically significant only in the HeH group (HeH *P*=0.024 and *P*=0.036, respectively; non-HeH: *P*=0.7114 and *P*=0.4615, respectively, Spearman’s correlation). No correlations were seen between the expression of CTCF, cohesin-, or condensin-related genes and PCG percentage and CMS apart from between the levels of PCG and *SMC4* and *PDS5B* expression (*P*=0.027 and *P*=0.022, respectively, Spearman’s correlation) (**Supplementary Table 7**).

Thus, we find that aneuploidy in the form of chromosomal gains is associated with increased chromatin interactions across TAD boundaries, likely due to low CTCF levels, and genome-wide transcriptional dysregulation in BCP ALL, as well as with sister chromatid cohesion defects and poor metaphase chromosome morphology.

### Cis-regulatory architecture in BCP ALL and the ALL Micro-C database

Our Micro-C data analysis provided the opportunity to study chromatin loop interactions on a genome-wide scale, revealing the details of gene regulation to the highest resolution achieved so far in primary hematopoietic cells. By merging the data of the 35 BCP ALL cases, we reached a 1 kb resolution and identified 55,240 high confidence chromatin loops with a median span of 163 kb (range 5 kb-1.99 Mb; **Supplementary table 8**). Comparing with publicly available data on histone marks and chromatin accessibility from primary BCP ALL,^12,13^ the majority of loops had at least one anchor that overlapped with histone marker (n=49,491; 89.6%) and ATAC-seq peaks (n=52,697; 95.4%). We further classified the 55,240 loops as promoter-promoter (P-P) and enhancer-promoter (E-P) based on overlapping transcription start sites, resulting in 11,360 (21%) P-P loops and 18,220 (33%) E-P loops while the remaining were annotated as structural loops. Promoter interacting anchors were enriched for the enhancer-associated marks H3K4me1 and H3K27ac (*P*<2.2e-16, Fisher’s exact test). H3K4me1 was more prevalent, marking 87% of enhancer anchors, while H3K27ac covered 73% (**Figure 3A**). To assign chromatin loops to the regulation of specific genes, we utilized matched RNA-seq data, leading to the identification of regulatory loops targeting 15,093 transcripts from 10,863 genes. A total of 11,230 (74%) of these transcripts were found to be expressed (RSEM≥10) in at least half of the samples and annotated as such for the downstream analysis.

**Figure 3.**
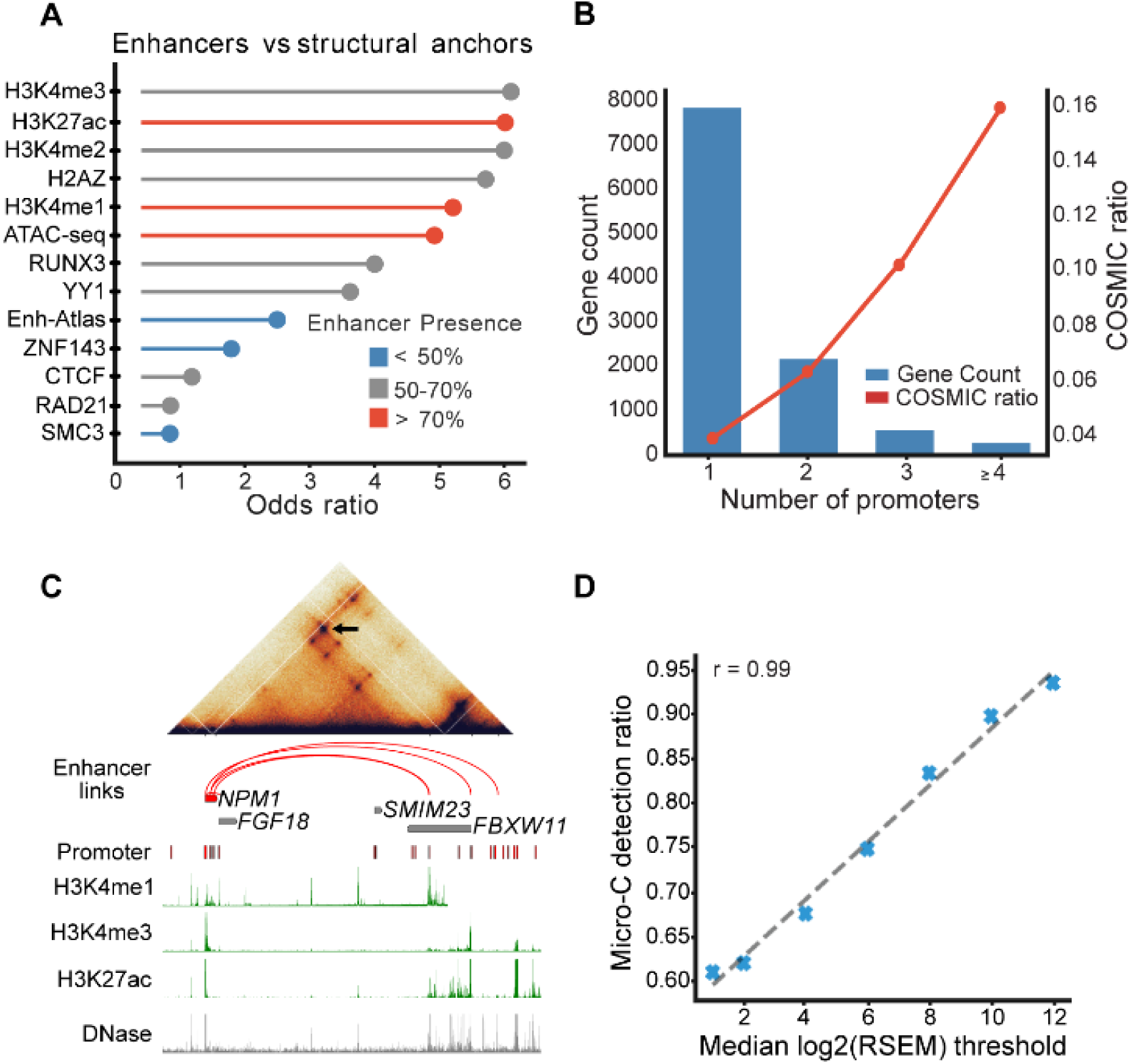
Overview of findings from 1 kb loop data regarding the cis-regulatory architecture. **A.** Lollipop plot showing the enrichment of selected marks on enhancer anchors compared with structural anchors. The x-axis shows odds ratios generated from Fisher’s exact test. **B.** Bar plot demonstrating the association between COSMIC gene presence and the number of identified promoters with Micro-C analysis. The x-axis shows the promoter count, y-axis shows gene count and the ratio of COSMIC genes on left and right sides respectively. Bars represent the gene count matched with the given number of promoters and the line shows the corresponding cosmic ratio. **C.** An example output from the database app showing the result of a query for *NPM1,* along with a heatmap generated from the region on top. The gene of interest is color coded in red and the arcs represent loops interacting with reported promoters of *NPM1*. D. The detection accuracy of expressed gene promoters by Micro-C with different log2(RSEM+1) cut-off values. A linear regression model shows a strong correlation coefficient of 0.995.

Analysis of histone marks for these loop anchors showed enrichment of the enhancer-associated marks H3K4me1, H3K4me3, and H3K27ac (*P*<2.2e-16, Fisher’s exact test), respectively, whereas transcripts with detectable chromatin loops but no detectable expression (n=3,863; 26%) were enriched for the repressive histone mark H3K27me3 (*P*=1.4e-54, Fisher’s exact test) (**Supplementary figure 6**). Finally, to evaluate these Micro-C-derived annotations against existing datasets, we compared our promoter-interacting anchors (n=32,077) with GM12878 enhancers from EnhancerAtlas^14^, finding an enrichment with an odds ratio of 2.41 (P<2.2e-16, Fisher’s exact test) and supporting the biological significance of the extracted loops. For enhancers, 15,204 (47%) of the annotations overlapped with those in the database while the rest remained as new findings, showing the strength of Micro-C for cis-regulatory elements (CRE) discovery. Taken together, we hereby provide a genome-wide map of CREs affecting the expression of more than 10,000 genes in BCP ALL.

Next, we examined the promoter utilization of genes found within our interaction network. For most genes with detected chromatin interactions (n=7,850; 72.3%), a single unique promoter was identified. However, 3,013 genes utilized multiple promoters (median 2, range 2-9). To gain insights into their potential role in leukemogenesis, we performed over representation analysis, revealing significant enrichment of several leukemia-associated pathways, including the Rho GTPase cycle, MAPK activation, and BCR signaling (**Supplementary table 9**). Moreover, genes with multiple promoter usage were significantly enriched in the COSMIC cancer gene census^15^ (*P*=8.55e-17; Fisher’s exact test), with an odds ratio of 2.14 and a general trend towards increased usage of different promoters (**Figure 3B**). For example, *NPM1*, one of the most frequently targeted genes in AML, has previously been reported to have increased levels of alternative transcripts in both AML and ALL compared to healthy samples^16^. Here, we found six chromatin loops targeting four promoters of *NPM1* (**Figure 3C**), two of which belong to transcripts that were expressed in at least half of the samples in our dataset, revealing the CREs underlying these transcripts.

We also explored the extent to which the strength of promoter detection in Micro-C is associated with expression. Higher expression levels were observed for genes with loop interactions. Furthermore, Micro-C detection sensitivity also correlated strongly with expression, showing a correlation coefficient of 0.995 in a linear model between RSEM threshold and loop presence (**Figure 3D**). For example, transcripts with median log2(RSEM+1) values greater than 12 were detected by Micro-C with 94% accuracy. Taken together, we demonstrate that alternative promoter usage is present in BCP ALL, in line with transcriptomic studies that have shown alternative promoter usage to be a critical mechanism underlying the majority of isoform switching events observed in various cancers^17^, strongly indicating that Micro-C can be used as a stand-alone tool to identify these events.

Taken together, our analysis provides detailed information of the 3D genome that forms a complex, multi-layered network with a vast amount of biologically relevant information yet to be uncovered. Key clinical features in BCP ALL such as recurrent mutations, structural variants, or subtype-specific gene expression patterns all operate within this dynamic regulatory architecture. To facilitate future research in this area, we developed the ALL Micro-C database that enables intuitive exploration of BCP ALL chromatin loops experimentally identified by our Micro-C analysis. Notably, since many of these regulatory loops are likely to be shared between similar cells, this database can also be utilized for exploring gene regulation in other hematopoietic malignancies and during normal hematopoiesis. The application provides a user-friendly interface to visualize loop structures across different genomic resolutions (1 kb and 10 kb bins) and includes a subtype-specific 10 kb module. Using this platform, researchers can 1) identify canonical and non-canonical promoter interactions for genes of interest, 2) visualize experimentally identified chromatin interactions at specific genomic loci, 3) compare interactions across BCP ALL subtypes to detect unique regulatory patterns, and 4) find E-P, P-P, and structural loops derived from Micro-C experiments based on customized queries. Figure 3C shows an example output for an *NPM1* query, showing its putative enhancers identified in 1 kb loop data from the merged cases. The application together with its code and documentation is available at https://github.com/efeaydin-bio/microc-all-db.

### Differential chromatin loops reveal subtype-specific regulation of leukemia-related genes

The above genome-wide identification of chromatin loops that regulate gene expression also allows exploration of specific CREs that underlies differences in gene expression. To address this, we aggregated the original 1 kb chromatin loop calling data into 10 kb resolution bins, to allow investigation of individual cases, and integrated with RNA-seq data. This revealed that alterations in chromatin interactions can account for previously reported differences in the expression of key leukemia-related genes among BCP ALL subtypes and provide deeper understanding of the molecular regulation of gene expression, providing novel insight into leukemogenesis. Notable examples of this included *HOXA9*, *FLT3*, *TP53*, *CD44*, *IKZF1*, *ERG*, and *XBP1* (**Figure 4A**), detailed as follows.

**Figure 4.**
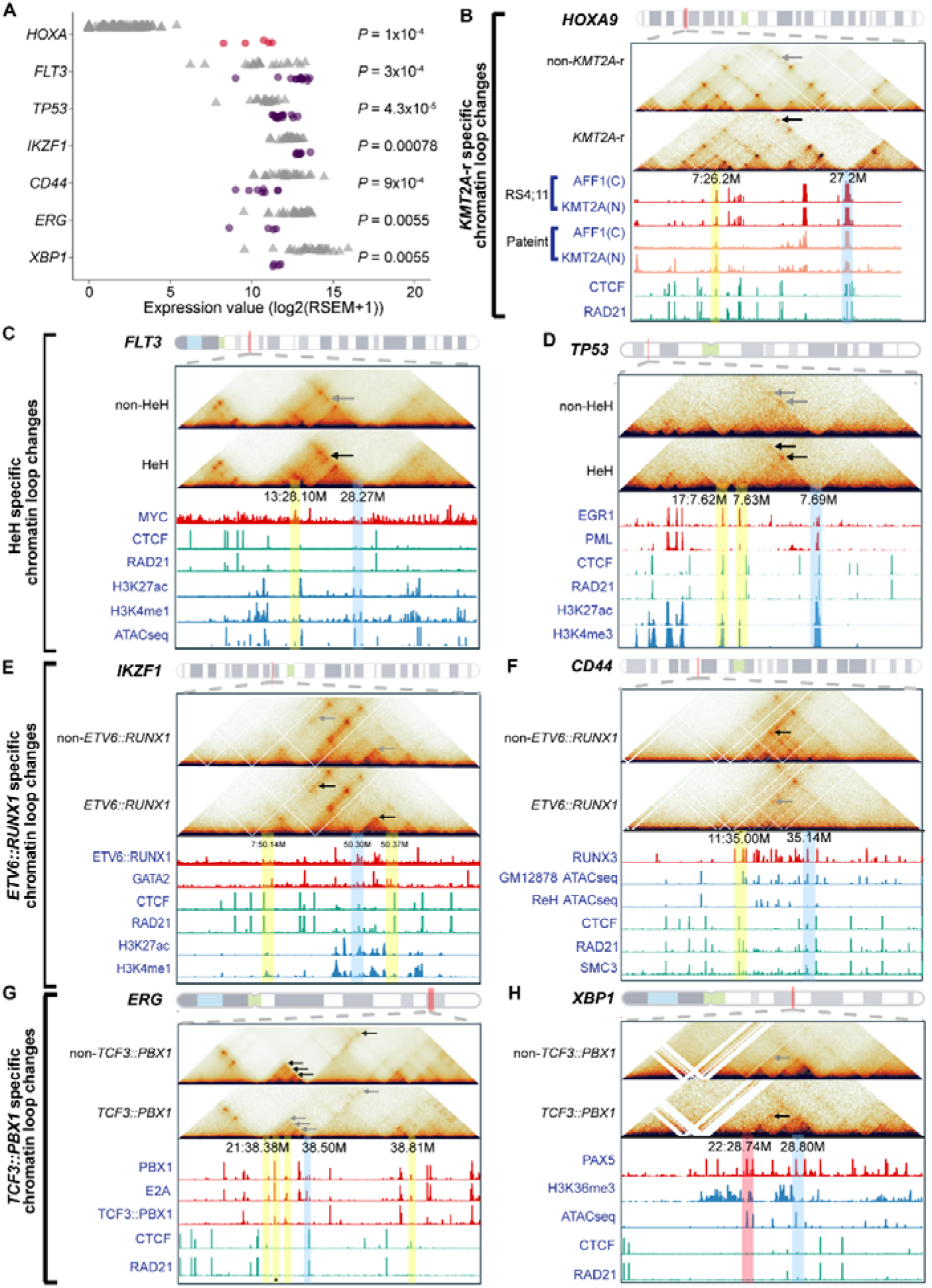
Chromatin organization and gene expression in B-cell precursor acute lymphoblastic leukemia (BCP ALL). **A.** Differential gene expression analysis of *HOXA3*, *HOXA6*, *HOXA7*, *HOXA9* and *HOXA10* (red) as well as *FLT3*, *TP53, IKZF1*, *CD44*, *ERG* and *XBP1* (purple) between cases from subtypes with altered chromatin loop strength and those without (grey). Panels **B-H** depict subtype-specific looping changes at the (B) *HOXA* cluster, (C) *FLT3*, (D) *TP53*, (E) *IKZF1*, (F) *CD44*, (G) *ERG* and (H) *XBP1* loci: the top row shows each locus’ genomic context on the reference chromosome; the middle row presents ICE-normalized, pooled Micro-C contact heatmaps comparing non-*KMT2A*-r vs. *KMT2A*-r (B), non-HeH vs. HeH (C-D), non-*ETV6::RUNX1* vs. *ETV6::RUNX1* (E-F) and non-*TCF3::PBX1* vs. *TCF3::PBX1* (G-H); and the bottom panels overlay ChIP-seq tracks, with red indicating transcription factor occupancy, green representing the CTCF/cohesin complex, and blue marking histone modifications. Arrows highlight subtype-specific loop strength alterations; gene promoters are marked by blue squares, enhancers by yellow squares and the *XBP1* silencer by a red square. The reported *ERG* risk locus rs2836371 is marked by an asterisk.

*HOXA* genes, particularly *HOXA9*, are uniquely overexpressed in the *KMT2A*-r leukemia^18,19^. We identified a chromatin loop specific to *KMT2A*-rearranged (*KMT2A*-r) case, with one anchor situated within the *HOXA* gene cluster, in close proximity to the promoters of *HOXA3*, *6*, *7*, *9*, and *10* genes. These genes are known to be highly expressed in *KMT2A*-r leukemias ^20^ and our study confirms their elevated expression in the investigated case (**Figure 4A**). Micro-C strongly suggests the presence of an enhancer-promoter interaction that is likely driving elevated *HOXA9* expression (**Figure 4B**). Notably, this loop is absent in other BCP ALL subtypes, suggesting that its formation is specific to the *KMT2A*-r subtype and that additional transcriptional regulators may be involved. To investigate further the underlying mechanism, we analyzed ChIP-seq and CUT&RUN data from primary patient samples and cell lines (including SEM and RS4_11) harboring the *KMT2A*::*AFF1* fusion^21,22^, revealing distinct peaks for KMT2A(N) and AFF1(C) at both anchors, which serve as defining markers of the *KMT2A*::*AFF1* fusion gene (**Figure 4B**). In contrast, although peaks for DOT1L, MLLT1, USF2, PSIP1, and MAZ were detected, these factors are not significantly upregulated in *KMT2A*-r cases, indicating that the *KMT2A*::*AFF1* fusion may primarily be responsible for establishing the enhancer-promoter loop, facilitating the recruitment of a specialized set of co-regulators that activate *HOXA* genes expression (**Figure 4B**). Moreover, the absence of MEN1 and MYB binding at the novel enhancer suggests that *HOXA* genes activation in these cases may be driven by an alternative regulatory mechanism, potentially mediated by the unique activity of the *KMT2A*::*AFF1* fusion rather than by the conventional MEN1 and MYB pathways.

Previous studies have shown that *FLT3* is intrinsically overexpressed at both the mRNA and protein levels in HeH and *KMT2A-r* BCP ALL, even in the absence of *FLT3* mutations^9,23^. Our prior work demonstrated that *FLT3* expression can be modulated by the two distinct enhancers DS2 and DS3 in HeH^24^. Here, we observed a marked increase in chromatin loop interaction strength between the *FLT3* promoter and DS3 in HeH and *KMT2A*-r, suggesting that this strengthened interaction underpins the high expression of *FLT3*. ChIP-seq data from the GM12878 cell line indicate that the *FLT3* promoter-DS3 interaction occurs independently of the CTCF/cohesin complex, implying that transcription factors are primarily responsible for stabilizing this loop. Indeed, both the *FLT3* promoter and DS3 show enrichment for key transcription factors, including MYC (**Figure 4C; Supplementary figure 7A**), and MYC ChIP-seq data from *KMT2A::AFF1* cell lines reveal a distinct binding peak at both the *FLT3* promoter and the DS3 enhancer region^25^. Given that *MYC* is known to have a consistently high expression level in HeH and *KMT2A*-r, it may thus play a pivotal role in driving the enhancer-promoter interaction at DS3, driving elevated *FLT3* expression and leukemogenesis.

*TP53* is one of the most important tumor suppressor genes^26^. Our analysis identified seven E-P loops at the *TP53* locus, with two loops exhibiting significantly enhanced interaction strength in HeH, associated with a higher *TP53* expression in this subtype. Transcription factor binding analysis revealed that four loops possessed robust EGR1 and PML binding at their anchors (**Figure 4D**). Both EGR1 and PML have been shown to directly regulate *TP53* expression in leukemia^27,28^, and our RNA-seq data confirm that they are highly expressed in HeH cases (**Supplementary figure 7B, C**). Thus, these transcription factors likely reinforce loop stability, facilitating the elevated *TP53* expression observed in HeH subtype. The effect of the relatively high *TP53* expression in HeH remains unclear, but we note that recent data suggest that these leukemias are genomically stable despite their aneuploidy^29^.

We also observed that *IKZF1,* a master transcription factor in hematopoiesis that is frequently targeted by deletions and mutations in BCP ALL^30^, was relatively highly expressed in *ETV6*::*RUNX1* cases, whereas HeH cases displayed markedly lower levels (**Figure 4A**). This differential expression appears to be partially driven by three distinct enhancer-promoter loops at the *IKZF1* locus (**Figure 4E**). For the first two of these, the interaction strength was substantially increased in *ETV6*::*RUNX1* samples and decreased in HeH cases. ChIP-seq data from the GM12878 cell line revealed clear binding peaks for CTCF and the cohesin subunit RAD21 at both anchors of this loop (**Figure 4E**). As CTCF and cohesin expression is significantly higher in *ETV6*::*RUNX1* than HeH^9^, this may account for the differences in loop interaction strength, with elevated levels reinforcing robust loop integrity in *ETV6*::*RUNX1* cases. Additionally, GATA2 binding peaks were present at both anchors of this loop in leukemia cell line ChIP-seq data^31^. Notably, previous functional studies demonstrated that *GATA2* knockdown results in reduced *IKZF1* expression^32^. Consistent with this, our RNA-seq data indicate that *GATA2* is significantly higher expressed in *ETV6*::*RUNX1* cases than in HeH cases, suggesting that *GATA2* may serve as a pivotal mediator of *IKZF1* activation (**Figure 4E, Supplementary figure 7D**). Finally, ChIP-seq data from the REH cell line show robust RUNX1 binding at both loop anchors and that the ETV6::RUNX1 fusion protein directly binds to the *IKZF1* promoter (**Figure 4E**)^33,34^. The second E-P loop, which also showed stronger chromatin interaction in the *ETV6*::*RUNX1* cases, is enriched for active histone modifications, including H3K4me3, H3K9ac, and H3K27ac. Although the precise function of it remains unclear, its presence further supports an active regulatory state at the *IKZF1* locus. Together, these observations suggest that the coordinated actions of CTCF/cohesin-mediated looping, RUNX1/ETV6::RUNX1 fusion protein activity, and GATA2 recruitment establish a robust regulatory architecture driving *IKZF1* expression in *ETV6*::*RUNX1* BCP ALL.

Previous studies have demonstrated that *ETV6::RUNX1*-positive cases have a characteristic CD27^pos^/CD44^low-neg^ immunophenotype^35^. Micro-C indicates that CD44 expression is maintained by an E-P loop in BCP ALL (**Figure 4F**) and ChIP-seq data from the GM12878 cell line showed that this loop is anchored by strong binding of RUNX1, RUNX3, CTCF, RAD21, and SMC3, as well as associated with active histone marks. However, our Micro-C analysis shows that the enhancer shifts from the active A compartment to the repressive B compartment in *ETV6*::*RUNX1* BCP ALL. Since the B compartment is linked to more compact, silenced chromatin, this shift likely affects the function of the E-P loop. As a result, the promoter may have a reduced ability to recruit the necessary transcription factors and other regulatory proteins. In addition, we found that *RUNX3* expression is significantly lower in the *ETV6*::*RUNX1* subtype compared to the other BCP ALL cases (**Supplementary figure 7E**). Taken together, the lower level of *RUNX3*, combined with the compartment shift, likely weakens the E-P loop and lead to the downregulation of *CD44* expression, contributing to the characteristic immunophenotype observed in *ETV6*::*RUNX1* ALL.

In *TCF3*::*PBX1* cases, we found low expression of *ERG* and differential loop strengths (**Figure 4G**). *ERG* encodes an ETS transcription factor and is frequently deleted in *DUX4*-r BCP ALL^36,37^. Micro-C analysis revealed a complex network of enhancer–promoter interactions regulating *ERG* expression, involving six E-P loops of which four exhibit a significant loss of interaction strength in cases with *TCF3*::*PBX1* fusion. ChIP-seq data from the *TCF3*::*PBX1* BCP ALL cell line 697 show that both the TCF3 and PBX1 proteins bind at the *ERG* promoter, suggesting that they maintain loop stability and active transcription^38,39^. Interestingly, however, the TCF3::PBX1 fusion protein does not^38^. This loss of binding indicates that the fusion protein may be unable to substitute for intact TCF3 and PBX1, leading to a disruption in enhancer-promoter communication. Further examination of the enhancer anchors of the affected loops revealed that three out of four contain binding peaks for both TCF3 and PBX1, while only two exhibit binding of the TCF3::PBX1 fusion protein (**Figure 4G**). Thus, the partial loss of intact TCF3 and PBX1, resulting from the t(1;19) that truncates these proteins, likely plays a crucial role in weakening these chromatin interactions and lowering *ERG* expression in *TCF3*::*PBX1*. Furthermore, prior genome-wide association studies have identified the *ERG* risk locus rs2836371 as being associated with *TCF3*::*PBX1* ALL^40^. Notably, one of the four weaker E-P loop anchors is located just 400 base pairs away from this susceptibility allele, implying that diminished loop strength in this region may further contribute to altered *ERG* regulation in some cases. Together, these findings suggest that intact *TCF3* and *PBX1* binding is crucial for sustaining robust enhancer–promoter interactions at the *ERG* locus. In *TCF3*::*PBX1* cases, the inability of the fusion protein to engage the promoter and the reduced binding at key enhancer anchors likely disrupts chromatin architecture and leads to the downregulation of *ERG* expression. Overall, our results suggest that *ERG* may have a previously unknown leukemogenic effect in *TCF3*::*PBX1* BCP ALL. Finally, *XBP1* expression was differentially regulated across BCP ALL subtypes, with significantly higher levels in HeH and lower in *TCF3*::*PBX1* (**Figure 4A**). XBP1 is a critical transcription factor involved in the unfolded protein response and B-cell differentiation, and its dysregulation has been implicated in leukemogenesis and cellular stress responses^41,42^. Our analysis identified a distinct chromatin loop at the *XBP1* locus (**Figure 4H**). In the *TCF3*::*PBX1* cases, this loop exhibited increased interaction strength. By screening the GM12878 ChIP-seq dataset, we detected clear PAX5 binding peaks at both anchors of the loop, along with prominent H3K36me3 signals—an epigenetic mark typically associated with heterochromatin formation and gene repression. This repressive chromatin environment is consistent with the observed downregulation of *XBP1* in *TCF3*::*PBX1* cases. Previous studies^42–44^ of B-cell lymphomas have demonstrated that PAX5 acts as a repressor of *XBP1* expression, establishing a functional link between PAX5 binding and the suppression of *XBP1* transcription. Although our RNA-seq analysis of the Micro-C cases did not reveal any statistically significant differences in *PAX5* expression among HeH, *TCF3*::*PBX1*, and other BCP ALL subtypes, the observed expression trends support the repression mechanism (**Supplementary figure 7F**). Moreover, analysis of a larger RNA-seq cohort^45^ supports the hypothesis of PAX5-mediated repression of *XBP1* in *TCF3::PBX1*-positive cases (**Supplementary figure 7F**). Taken together, our findings suggest that the PAX5-associated chromatin loop, enriched with repressive H3K36me3 marks, plays a key role in downregulating XBP1 in *TCF3*::*PBX1* cases. Finally, our data demonstrate that Micro-C can be effectively used to identify CREs that function as repressors, thereby providing novel insights into the epigenetic regulation of gene expression in leukemia.

The genes highlighted here constitute some of the most prominent driver genes in pediatric BCP ALL. Collectively, these findings provide a comprehensive view of how chromatin looping, transcription factor dynamics, and nuclear compartmentalization converge to modulate gene expression. Our Micro-C dataset provides an opportunity for detailed analyses of the regulation of many more genes.

### Structural rearrangements change chromatin interactions in BCP ALL

Structural variants (SVs) lead to changed chromatin interactions, both as a direct effect of rearranging the genome and indirectly in the form of position effects^46,47^. Micro-C was used to identify SVs, detecting a total of 166 SVs with sizes ranging from 55 kb to 32 Mb, comprising 121 intrachromosomal rearrangements (deletions, duplications, and inversions), 16 translocations associated with the canonical genetic subtypes, and 29 additional translocations in 35 BCP ALL cases (**Supplementary table 11**). Supporting data were available for 33 of the included cases, derived from whole genome sequencing (WGS) (n=20), single nucleotide polymorphism (SNP) array (n=32) or single-cell (sc) WGS (n=8) data (**Supplementary table 1**). For cases with available WGS data, it was also possible to distinguish between somatic and germline variants. Among the 72 SVs validated with WGS data, 51 were somatic and 21 were germline variants. Detected somatic variants included well-known leukemia driver events with prognostic impact, such as focal deletions of *PAX5* and *ETV6* in addition to the subtype-defining translocations (**Supplementary figure 8; Supplementary table 11**). Notably, one of the validated events was a subclonal translocation between chromosomes 4 and 19 in HeH_6 that had been missed by bulk WGS but detected at a frequency of 2.8% of cells by scWGS, showing the potential of Micro-C to detect SVs present in very small subclones (**Figure 5A**). In total, 108/125 (88.64%) of the SVs detected by Micro-C were validated by supporting data, showing that Micro-C can be a powerful tool to investigate SVs.

**Figure 5.**
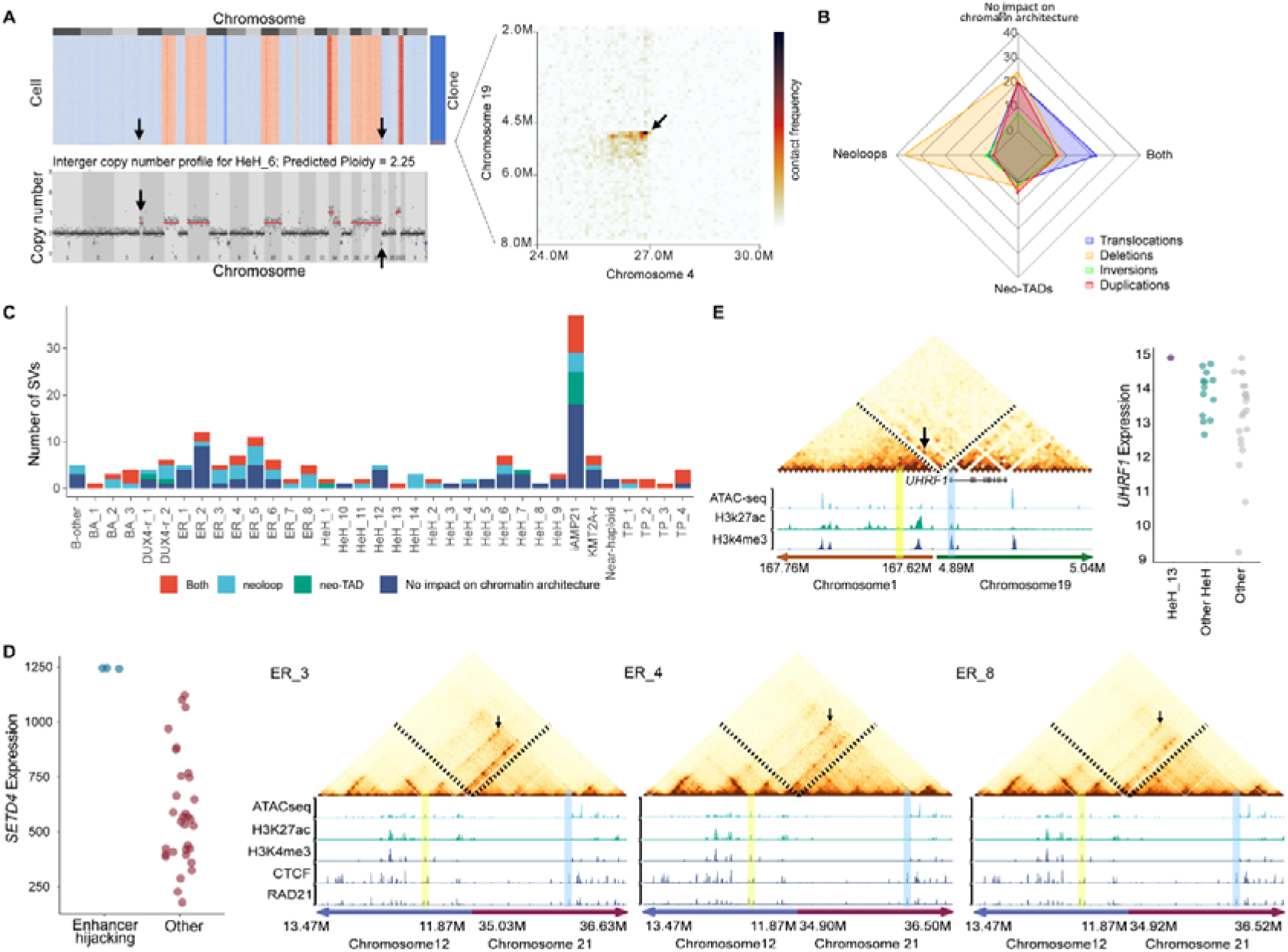
Structural variants (SVs), neo-topologically associating domains (TADs), and neoloop events in B-cell precursor acute lymphoblastic leukemia (BCP ALL). **A.** Detection of a subclonal translocation between chromosomes 4 and 19 in case HeH_6. ICE-normalized Micro⍰C contact maps (100 kb resolution, right panel) reveal a translocation event, which was independently identified by single-cell whole-genome sequencing (scWGS) at a frequency of 2.8% (left top). The copy number of individual leukemic blast cells (rows) are color coded, with red corresponding to chromosomal gains and clone identity shown on the right. The scWGS-based copy number profile shows breakpoints in chromosomes 4 and 19 (left bottom). **B.** Radar plot depicting the percentage of SVs that induce specific chromatin architectural alterations. Four SV types – translocations, deletions, inversions, and duplications – were evaluated along axes representing distinct outcomes: formation of a neoloop, formation of a neo-TAD, simultaneous formation of both a neoloop and a neo-TAD, or no detectable effect. Each vertex represents the number of the corresponding SV type that leads to the specified outcome. **C.** Bar plot summarizing the number of structural variants that resulted in neoloop and neo-TAD formation across BCP ALL cases. **D.** Cases ER_3, ER_4, and ER_8 exhibit neoloops resulting from a balanced t(12;21) translocation. The left panel shows *SETD4* gene expression in BCP ALL cases with and without an enhancer-hijacking event. The right panels show ICE-normalized Micro-C contact heatmaps that reveal distinct neoloop formations associated with recurrent t(12;21) translocations. In the heatmaps, black arrows highlight specific chromatin interactions between gene promoters (marked by blue rectangles) and the newly hijacked enhancers (marked by yellow rectangles). The bottom panels present ATAC-seq and ChIP-seq tracks for H3K27ac, H3K4me3, and the CTCF/cohesin complex (derived from the GM12878 cell line). **E.** ICE-normalized Micro⍰C contact heatmaps (left top panel) display a neoloop in case HeH_13 caused by a somatic t(1;19) translocation at 10 kb resolution. The black arrow highlights a chromatin interaction between the *UHRF1* promoter (blue rectangle) and an enhancer located on chromosome 1 (yellow rectangle), indicative of a potential enhancer-hijacking event that results in *UHRF1* upregulation. The middle panels show ATAC-seq tracks and ChIP-seq tracks for H3K27ac and H3K4me3 from the GM12878 cell line. The right panel compares *UHRF1* expression levels in case HeH_13, other HeH cases, and non-HeH cases.

To look at position effects, i.e., how rearrangements of the genome result in changes in gene expression, we examined the impact of the recurrent SVs on TAD boundary alterations (neo-TADs) and chromatin loops (neoloops). Analysis by SV type revealed that translocations altered looping or domain structure in over half of cases: about 50% of translocations produced both neoloops and neo-TADs. Deletions disrupted organization in two-thirds of cases, with 66% forming neoloops and 15% establishing neo-TADs. Inversions were detected in 23 % of BCP-ALL cases (8/35); of these, 45% gave rise to neo-TADs and 50 % to neoloops. By contrast, duplications had smaller effects; 31 % induced neo-TADs and 17 % neoloops, while 66% showed no impact on chromatin architecture. Overall, SVs thus had a large impact on local chromatin interactions, with translocations and deletions being most disruptive (**Figure 5B**).

We identified 64 neo-TADs caused by 50 somatic SVs and 3 caused by 3 germline SVs (range 0-24, median 1; **Figure 5C, Supplementary Table 12**). These included 12 deletion-, 8 inversion-, 10 duplication- and 36 translocation-induced neo-TADs. Over 80% of the neo-TAD boundaries were enriched for CTCF ChIP-seq peaks and cohesin complex ChIP-seq peaks, suggesting that their formation was mediated by a CTCF/cohesin-dependent mechanism. The major subtype-defining translocations t(1;19), t(4;11) (*KMT2A*-r), t(9;22), and t(12;21) all resulted in neo-TADs at the translocation breakpoints, indicating additional effects apart from the formation of fusion genes. Notably, we observed neo-TADs caused by (12;21)(p13;q22) in five *ETV6*::*RUNX1* cases with a balanced t(12;21) translocation, whereas no discernible neo-TADs in the same regions were detected in three *ETV6*::*RUNX1* cases with more complex rearrangements, suggesting different chromatin reorganization in these two genetic types of *ETV6*::*RUNX1*. The iAMP21 case, characterized by intrachromosomal amplification of chromosome 21, showed 20 neo-TADs that were caused by 15 SVs in this chromosome, highlighting the extensive genomic reorganization that disrupts the native chromatin landscape in this BCP ALL subtype, which may in turn may create novel regulatory circuits and contribute to the dysregulation of gene expression. In contrast, no neo-TADs were detected in the *DUX4*-r cases, likely due to the translocation occurring in repetitive regions that hinder reliable neo-TAD analysis.

We found 735 neoloops generated by 88 SVs across the 35 BCP ALL cases (range 0-135 per case, median 9), 53% of which were associated with neo-TAD-forming SVs (**Figure 5C; Supplementary Table 12**). Eleven genes were targeted by neoloops in enhancer-hijacking events associated with their increased expression. Recurrent overexpression of *SETD4* (in chromosome 21), encoding a SET domain methyltransferase that has been implicated in hematopoiesis and stem cell regulation^48^, was seen in 3/5 *ETV6*::*RUNX1* cases harboring a balanced t(12;21), which also had the neo-TAD described above. Further investigation of the neoloop anchors in these cases revealed a potential enhancer within intron 2 of *MANSC1*, approximately 500 kb distal to *ETV6* in chromosome 12, suggesting a previously unknown mechanism for *SETD4* activation in some cases from this genetic subtype (**Figure 5D**). We also observed recurrent overexpression of *LRP6*, a low-density lipoprotein receptor in 12p13.2 involved in the WNT pathway and implicated in AML^49^, which resulted from two distinct enhancer-hijacking events: one in a *DUX4*-r case with a focal *ETV6* deletion and another in an *ETV6*::*RUNX1* case. In the *DUX4*-r case, a neoloop-driven chromatin interaction was identified between the *LRP6* promoter and CREs located 1.5 Mb distal to *LRP6* in chromosome 12, whereas in the *ETV6*::*RUNX1* case, a novel enhancer in chromosome 21 interacted with *LRP6* across the translocation breakpoint (**Supplementary figure 9A**). Additionally, two enhancer-hijacking events were identified as resulting from the t(1;19) in two *TCF3*::*PBX1* cases, where the gene *REXO1* in chromosome 19 was activated by enhancers within *PBX1* intron 2 in chromosome 1 (**Supplementary figure 9B**). Furthermore, we observed an enhancer hijacking event driven by an unbalanced t(1;19) translocation (not resulting in the *TCF3*::*PBX1* fusion) in case HeH_13. This rearrangement resulted in the loss of part of one copy of *UHRF1*, an oncogene overexpressed in various cancers and involved in apoptosis inhibition by inducing epigenetic silencing of tumor suppressor genes^50,51^. ChIP-seq data from the GM12878 cell line reveal that both the *UHRF1* promoter and the novel enhancer, located in intron 1 of *RCSD1*, are enriched for RUNX3 binding and activating histone marks, including H3K27ac, H3K36me3, H3K79me2, and H3K4me1/2 (**Figure 5E**). Taken together, these findings suggest position effects and show that SV events may dysregulate gene expression via enhancer hijacking, revealing novel mechanisms that likely contribute to the development of BCP ALL.

## Discussion

Chromatin architecture is a major player in gene regulation^2^. Here, we present the first complete map of the 3D chromatin landscape in pediatric BCP ALL, including all major genetic subtypes and with a combined resolution of 1 kb. This gives unprecedented insights into chromatin organization and gene regulation, not only in leukemia but also in hematopoietic cells in general, considering the scarceness of high resolution Hi-C data from primary hematopoietic tissue^4,9,24,52–54^.

In terms of overall chromatin organization, our investigation showed that BCP ALL is more similar to hematopoietic stem and precursor cells than to more mature hematopoietic cells, in line with the immature phenotype of the leukemic cells. Furthermore, on all different levels, we found that somatic genetic aberrations were associated with specific chromatin states, from compartmentalization of the genome to TADs and enhancer-promoter interactions. This comprised not only the impact of established genetic subtypes but also position effects resulting from structural rearrangements. A notable finding was that all cases with a hyperdiploid chromosome number, including those belonging to other genetic subtypes than HeH (defined as 51-67 chromosomes without any disease-specifying gene fusions or evidence of duplicated near-haploid/low hypodiploid clones), displayed similar TAD structure. Thus, aneuploidy appears to have a major impact on TAD organization that even overrides a strong driver fusion gene such as *BCR*::*ABL1*. Notably, TAD boundary strengths were generally reduced in aneuploid cases and associated with a global dysregulation of gene expression that was not specific for the extra chromosomes. We have previously reported that low expression of CTCF at both the mRNA and the protein levels is a general feature of HeH^9^. Since CTCF binding is one of the main determinants for TAD boundary formation, this could explain the reduced TAD strength in the HeH cases. In line with this, analysis of the number of CTCF binding sites at TAD boundaries in relation to boundary loss or gain indicated an association between low *CTCF* and TAD boundary loss in HeH BCP ALL. Taken together, this study underscores that the HeH genetic subtype specifically, and possibly BCP ALL with hyperdiploid chromosome numbers generally, have aberrant TAD structure associated with gene dysregulation. An intriguing possibility is that this is associated with the hyperdiploidy itself, since the larger genome size presumably leads to a shortage of architectural proteins in the cell, at least for *CTCF*, which is encoded in chromosome 16; a chromosome that is rarely gained in HeH^29^. Further studies are needed to elucidate the role of this in leukemogenesis.

We identified >17,000 novel regulatory elements, highlighting the novelty and strength of applying Micro-C to primary patient samples for discovery of regulatory elements that may be missed in cell lines. Analyzing these data, we found regulatory loops underlying gene expression differences between different BCP ALL genetic subtypes for the known drivers *HOXA9*, *FLT3*, *TP53*, *CD44*, *IKZF1*, *ERG*, and *XBP1* in the absence of somatic aberrations targeting these genes. For example, although *ERG* is known to be involved in the leukemogenesis of primarily *DUX4*-r BCP ALL via targeted deletions, we here show that it is also differentially regulated via chromatin looping, and thereby implicated, in *TCF3*::*PBX1*-positive cases without any accompanying somatic genetic abnormalities. To promote further research, we provide an accessible web application summarizing the chromatin interaction landscape in BCP-ALL. Overall, our analysis is a step forward in understanding the regulatory network of genes that are involved in leukemogenesis.

In summary, we here report a thorough, high-resolution investigation of chromatin 3D structure in primary BCP ALL, the most common pediatric malignancy. Our study provides important data on how chromatin architecture regulates gene expression at different levels and offers novel insights into the role of chromatin 3D structure in leukemogenesis.

## Methods

### Patients and samples

Based on availability, either viable cells (n=32) or frozen cell pellets (n=5) from 37 bone marrow samples (**Supplementary table 1**) acquired at diagnosis, or during remission follow-up, at Skåne University Hospital, Lund, Sweden, were selected for Micro-C. The cohort comprised fourteen HeH, eight *ETV6*::*RUNX1*, four *TCF3*::*PBX1*, three *BCR*::*ABL1*, two *DUX4*-rearranged, one near-haploid, one *KMT2A*-rearranged, one iAMP21, and one B-other pediatric BCP ALL, in addition to one MPAL and one remission sample. Written informed consent was obtained from patients and/or guardians according to the Declaration of Helsinki and the study was approved by the Swedish Ethical Review Authority (Dnr 2023-01550-01). For normal hematopoietic cells, B-cell progenitors were sorted by fluorescence activated cell sorting from pools of frozen cord-blood. Cells were thawed and directly stained for surface antigens CD34, CD38, CD10, CD19 and CD20. All cells included within the CD34+CD38low/+ gate co-expressing CD10 and CD19 were sorted as pro-B cells, while CD34+ CD38low/+ cells single positive for CD19 were sorted as pre-B cells. CD20 expression was used to confirm its gradual acquisition along the maturation of the progenitors. Sorted cells were subsequently centrifuged and the cell pellets were snap-frozen in dry ice awaiting further processing. For Micro-C, multiple sorts of pro-B and pre-B cells were combined.

### Micro-C

Samples were prepared using the Dovetail Micro-C kit following the manufacturer’s protocol, with adjustments in enzyme and fixative concentration depending on the number of input cells. Briefly, in-nucleus fixation with disuccinimidyl glutarate (DSG) and formaldehyde was carried out, followed by digestion with micrococcal nuclease. Cells were lysed using sodium dodecyl sulfate (SDS) and chromatin capture beads were used to rescue the cross-linked chromatin fragments, followed by proximity ligation and crosslink reversal. Next, DNA was purified and libraries were constructed with Illumina-compatible adaptors. High-throughput pair-end sequencing was performed by the Center for Translational Genomics (CTG), Lund University, using an Illumina NovaSeq 6000 platform, to generate on average 3.1 billion 2 x 150 bp read pairs.

### RNA-seq library preparation, data processing, and differential gene expression analysis

RNA-seq libraries were prepared with the Illumina Stranded mRNA Prep workflow (Illumina, San Diego, CA). Paired-end reads of 36 patients and one healthy control were generated with an average depth of 30 million read pairs (2×150bp). Paired-end reads were aligned to the hg38 genome with the STAR 2-pass mapping pipeline^55^. Read counts for protein-coding genes and their corresponding transcripts were quantified using RSEM^56^. Genes with RSEM values less than 10 in 50% of the samples were excluded. Benjamini–Hochberg-adjusted (BH-adjusted) Mann-Whitney U test *P* < 0.1 were applied to identify the differentially expressed genes.

### Structural variants analysis of SNP data and WGS data

Log R ratio (LRR) and B allele frequency (BAF) of SNP array data from Illumina (.idat files) and Affymetrix (.CEL files) intensity files were analyzed by Illumina GenomeStudio (v2.0, Illumina, San Diego, CA) and Affymetrix Analysis Power Tools (v2.10.0, Thermo Fisher Scientific Inc., Waltham, MA), respectively. Copy number alterations were called using TAPS and manually reviewed in GenomeStudio or Chromosome Analysis Suite (v3.3, Thermo Fisher Scientific Inc., Waltham, MA). In the case of Illumina platform-generated sequencing data, paired-end reads alignment was executed using the GDC DNA-Seq analysis pipeline (https://github.com/NCI-GDC/gdc-workflow-overview?tab=readme-ov-file) and structural variants were called using Manta^57^ and Gridss2^58^.

### Micro-C data processing

Micro-C sequencing data were processed using the 4DN Hi-C data processing pipeline (https://data.4dnucleome.org/resources/data-analysis/hi_c-processing-pipeline). Pairtools was used to sort, remove duplicate reads and extract uniquely mapped contacts. Micro-C contact matrices were generated at multiple resolutions (range from 5 kb to 1 Mb) using Juicertools^59^ and Cooler^60^. Normalization of contact matrices was performed using Juicertools “scale”, and caICB^61^ algorithms.

### Compartment analysis

FanC^62^ was used to define compartments from all Micro-C samples using the Juicer tools SCALE algorithm normalized 500 kb resolution contact matrices as input. Since the compartments eigenvector positive/negative values are initially arbitrary, the ‘A’ (active) and ‘B’ (inactive) compartments were defined using ATAC-seq data from the REH cell line and primary HeH cases (unpublished data). ATAC-seq sequencing reads were aligned to human reference genome build hg38 using bwa and peak calling was performed by Genrich (https://github.com/jsh58/Genrich). The number of peaks in each 500 kb bin was calculated using BEDtools^63^ module map. Subsequently, bins for each chromosome were segregated into two groups based on the FanC-generated compartments eigenvector value (>0 or <0). Bins with eigenvector values equal to zero represent ambiguous regions and were therefore excluded from further analysis. The Mann-Whitney U test was then applied to identify the group with a significantly higher peak number and the bins in that group were defined as A (active) compartment. The 500 kb bins were used to ascertain A/B compartment shift events in different groups. Publicly available Hi-C data from seven healthy samples, of which four were peripheral blood mononuclear cell samples and three CD34+ hematopoietic stem cell-enriched samples^4^, were also included in the open chromatin comparison. Regarding compartment shifts analysis, we considered it a shift when every case from a group was in a different chromatin state than the comparing group. TFBSTools^64^ and JASPAR2022^65^ bioconductor packages were used to find binding motifs on selected compartments. Position frequency matrices for RUNX1 (MA0002.1) and PBX1 (MA0070.1) were retrieved, and corresponding position weight matrices were constructed using a log2 probability ratio. Motif occurrences were identified using a minimum score threshold of 85% relative to the maximum possible score.

### TAD boundary analysis

The SCALE algorithm corrected 25 kb contact matrix was utilized to predict TAD structures using the Juicer tools module arrowhead for each individual sample. TAD boundaries detected across different samples were merged, removing redundancies by integrating those within a ±50 kb window. The merged TAD boundaries were then used to calculate boundary scores for each sample with Chromosight^66^. Principal component analysis was performed based on the informative boundary scores called in all samples. To compare TAD boundary score changes between different groups, the Mann-Whitney U test was performed on the boundary scores data, using a significance threshold of BH-adjusted *P* <= 0.01. To detect CTCF and RAD21 binding sites occupancy over the subgroup-specific boundaries, publicly available data from the GM12878 cell line were downloaded. CTCF and RAD21 chromatin immunoprecipitation sequencing (ChIP-seq) data were used to identify all CTCF/RAD21 peaks located with a ±25 kb tolerance around the boundary by the BEDTools^63^ module intersectBed.

### Chromatin loop analysis

Chromatin loop calling was performed at 1 kb resolution using Mustache^67^ on caICB-corrected pooled contact matrices from all the BCP ALL samples. To account for positional shifts introduced by loop-calling algorithms and the detection of neighboring chromatin sites of interacting regions due to close proximity, we extended each loop anchor by ±2 kb. We then removed loops spanning less than 5 kb genomic distance. To examine the epigenetic modifications, chromatin accessibility and transcription factors binding profile at loop anchors, histone marks (H3K4me1, n=8; H3K4me3, n=10; H3K27me3, n=4; H3K27ac, n=10) and ATAC-seq (n=156)^12,13,68^ were also incorporated. In addition, GM12878 transcription factors ChIP-seq data (n=78) from the ENCODE project were also incorporated^68^. To evaluate the presence of binding peaks for the factors described above at the loop anchors, intersections were performed between the peak regions and the loop anchor positions (defined at 1 kb resolution) with a ±2 kb tolerance to account for possible shifting biases. All transcription start sites (TSS) from protein coding transcripts in ENSEMBL release 113 (https://www.ensembl.org/index.html) were downloaded and assigned to their respective anchors. Enhancers previously identified in GM12878 were downloaded from Enhancer Atlas^14,69^. Based on the existence of TSS and reported enhancers and enhancer-associated histone modifications (H3k4me1 / H3k27ac) from GM12878 chipseq data on one anchor or both anchors, loops were annotated as E-P or P-P loops respectively. The remaining loops were considered structural. Median RSEM values from the paired RNA-seq data were assigned to anchors with corresponding transcripts. If more than one transcript was present in an anchor, the transcript with the highest value was used^70–72^. To check for the efficiency of Micro-C in detecting alternative promoter interactions, we clustered transcripts with TSSs within 5 kb distance together to minimize artificial interactions among closely spaced transcripts and included the most expressed ones for analysis. Comparisons between different types of anchors were carried out using the Fisher’s exact test.

To examine the chromatin loop strength for each BCP ALL sample, Chromosight^66^ quantify was used to calculate loop interaction scores for each sample at a coarse resolution (10 kb). Only loops with informative strength scores in at least 50% of samples within the same subtype were retained. To compare loop strength changes between subtypes, the Mann-Whitney U test was performed on the loop strength score data, using a significance threshold of BH-adjusted *P* <=0.1.

### Database web application

All the loops from merged 1 kb data were acquired as described above. An additional lower resolution version of these were created by binning the anchors to 10 kb bins. Furthermore, 10 kb subtype specific loops from *ETV6*::*RUNX1*, HeH, *TCF3*::*PBX1*, *DUX4-r*, and *BCR*::*ABL1* cases were called using mustache algorithm with a *P*-value threshold of 0.01^67^. All .tsv outputs were joined together for a master dataset. These loops were annotated based on the presence of TSS on their anchors as described above. An additional promoter track to help visualization was created by extending TSSs of all coding genes to include 500 bp downstream and 2000 bp upstream windows. Additional ENCODE tracks were created based on the bigWig files of H3K4me1, H3K4me3, H3K27ac, H3K27me3, and DNase experiments derived from GM12878. A database app to represent these regulatory interactions within a graphical user interface together with the relevant epigenetic tracks was created using Streamlit (https://github.com/streamlit) and deployed to https://github.com/efeaydin-bio/microc-all-db. PyGenomeTracks module was used for track visualization^73^.

### Cytogenetic assays

Metaphase spreads from 36 samples of our cohort (all except HeH_2 due to lack of material for cytogenetic preparations), stored from 1 to 31 years in 3:1 methanol:acetic acid fixative, were included in this analysis. Data from six cases on cohesion defects and from three cases on CMS have been previously published^9,10^ (**Supplementary table 1**). Cytogenetic assays used both standard G-banding and fluorescence *in situ* hybridization (FISH) preparations. An average of 28 metaphase images were captured using a Z2 fluorescence microscope (Zeiss, Oberkochen, Germany) and the karyotyping software CytoVision (Leica, Wetzlar, Germany). Analysis of cohesion defects was performed as previously described^10^, with defects being defined as visible gaps between sister chromatids, i.e., presence of primary constriction gaps (PCGs), and severity classified based on the number of affected chromosomes per cell; levels of PCGs per case are shown as percentages. Chromosome morphology analysis was performed as previously described^9^, with each metaphase being scored from 1 to 3 (poor, fair, or good morphology) based on band resolution, condensation and appearance, and the chromosome morphology score (CMS) was calculated for each case as the average score of its cells. Statistical analysis of cytogenetic features (PCG percentage and CMS) versus mRNA expression (Log2(RSEM)) was carried out using Spearman’s correlation including all BCP ALL cases, only HeH and only non-HeH cases. The investigated genes were the condensin subunits *NCAPD2*, *NCAPD3*, *NCAPG*, *NCAPG2*, *NCAPH*, *NCAPH2*, *SMC2* and *SMC4*, and cohesin subunits *PDS5A*, *PDS5B*, *RAD21*, *SMC1A, SMC3*, *STAG1* and *STAG2*. Comparison between cytogenetic features versus number of TAD boundaries was also performed by Spearman’s correlation.

### Detection of structural variants

Micro-C contact maps of 35 of the primary BCP ALL cases were included in the SV detection, excluding the MPAL and remission samples. EagleC^74^ was used to identify SVs based on the Micro-C contact matrices at 50 kb, 10 kb, and 5 kb. All hits from EagleC were manually verified using Juicebox (version 1.11.08) and only visually confirmed SVs were included in this study. WGS, SNP array and single-cell WGS data from previous studies (**Supplementary table 1**) were used to validate the EagleC findings. Cancer-related and leukemia-related genes of interest affected by SVs were annotated, as well as de novo chromatin loops caused by these SVs. NeoLoopFinder^75^ modules neoloop and neotad callers were used to identify neoloops and neo-TADs using the EagleC calling result as input, respectively.

## Supporting information

Supplementary tables description

## Data availability

The Micro-C and RNA-seq data generated in this study have been deposited in the European Genome Archive (EGA) under accession numbers EGAS50000000764 and EGAS50000000763, respectively. These datasets are available under restricted access due to privacy concerns; access can be obtained for academic research by contacting the Data Access Committee via EGA. The GM12878 histone marks (H3K27ac, H3K4me1, and H3K4me3), CTCF/RAD21 ChIP-seq, and DNase-seq data (narrowPead file) were downloaded from the UCSC database under accession code (http://hgdownload.cse.ucsc.edu/goldenpath/hg19/encodeDCC/wgEncodeAwgTfbsUniform/). The REH cell line ATAC-seq data were downloaded from the NCBI SRA database under accession code SRR16677149. The Extended BCP-ALL RNA-seq dataset was obtained from the EGA under accession number EGAS00001001795.

## Acknowledgements

The authors would like to acknowledge Clinical Genomics Lund, SciLifeLab, and Center for Translational Genomics (CTG), Lund University, for providing expertise and service with sequencing and analysis. This study was supported by grants from the Swedish Childhood Cancer Foundation, grant numbers PR2020-0033 (MY), TJ2020-0024 (MY), PR2024-0058 (MY), PR2024-0002 (BJ), PR2018-0023 (KP), and PR2024-0033 (KP); the Swedish Cancer Fund, grant numbers 23 2694 PjF (BJ) and 22 2062 Pj (KP); Governmental funding of clinical research within the National Health Service, grant number ALFSKANE-623431 (KP); the Swedish Research Council, grant numbers 2020-01164 (BJ), 2020-00997 (KP), and 2024-02505 (KP); IngaBritt och Arne Lundbergs Forskningsstiftelse, grant number LU2019-0100 (KP), the Gunnar Nilsson Cancer Foundation (MY), and the Royal Physiographic Society of Lund (LHM-C and EW).

## Author contributions

LHM-C, MY and KP conceived the study; LHM-C, GTD, and ELW performed experiments; LHM-C, EA and MY analyzed data; AC, RG, LO-A, HL, TF, BJ, MJ, and AKH-A provided samples and clinical data and analyzed data; KP supervised the study; LHM-C, EA, MY and KP wrote the manuscript with input from all authors.

