## Supplementary figures for "The 3D genome of pediatric B-cell precursor acute lymphoblastic leukemia"

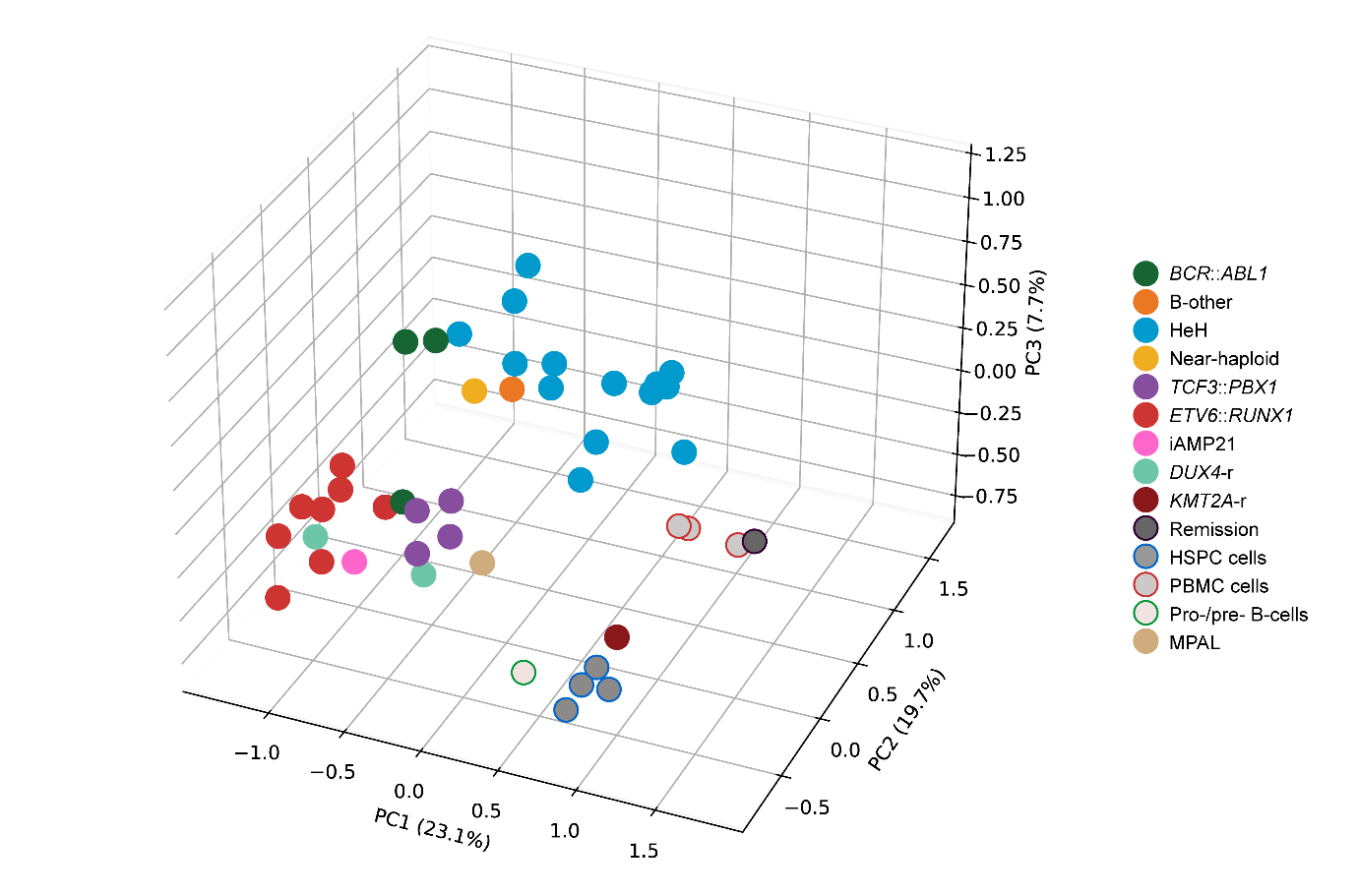

**Supplementary figure 1.** **Principal component analysis (PCA) of A/B compartments.** PCA was performed on the eigenvectors derived from Micro-C A/B compartment data calculated at 500 kb resolution. The PCA plot revealed clear clustering among the samples, with hematopoietic stem/progenitor cells (HSPC) and pro‑/pre‑B cells samples forming one distinct group, peripheral blood mononuclear cells (PBMC), the remission sample, and the *KMT2A*‑rearranged case clustering together in a separate group, and the remaining primary leukemia samples forming a broader, more diffuse cluster.

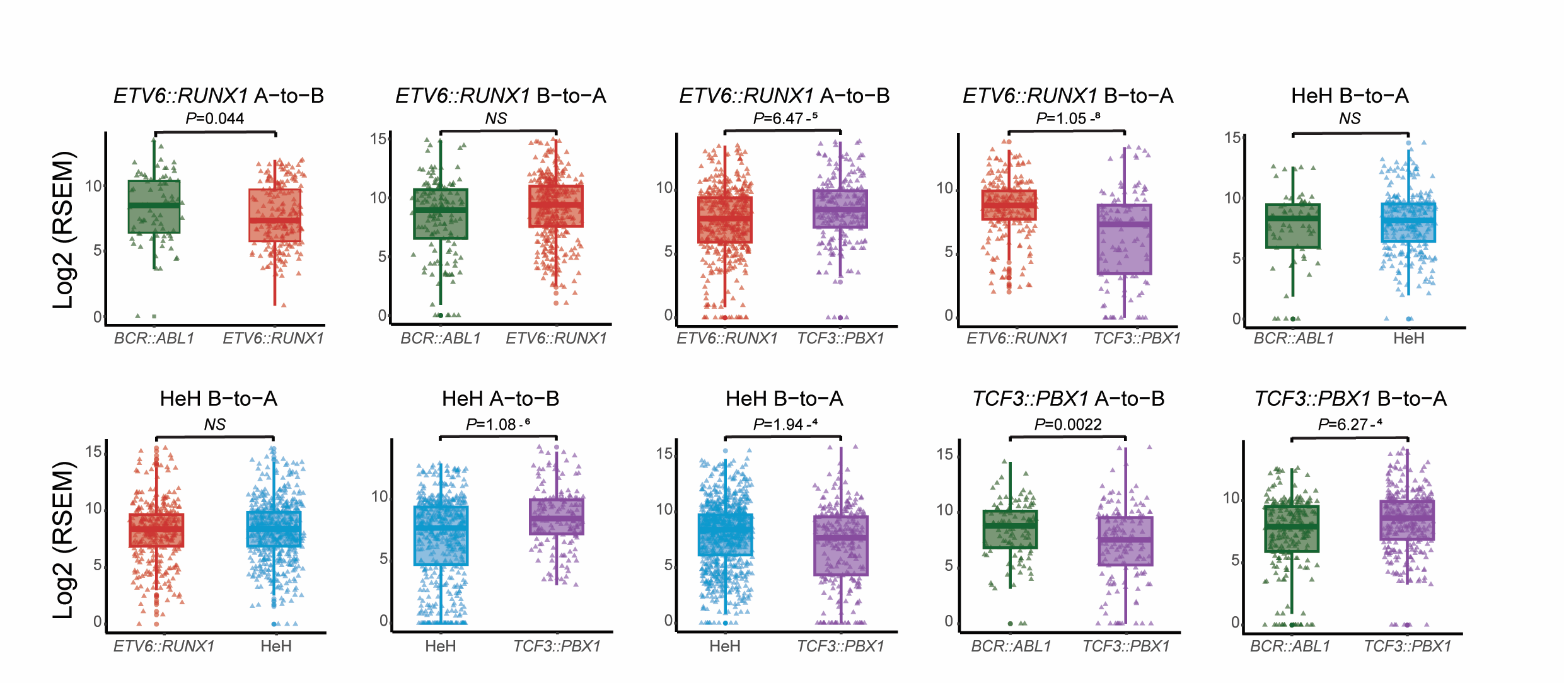

**Supplementary figure 2. Association between A/B compartment shifts, annotated from Micro-C data, and RNA sequencing data (mRNA levels displayed in Log2 RSEM).** Expression of genes located in B-to-A and A-to-B compartment shifts in *ETV6*::*RUNX1* against *TCF3*::*PBX1*, HeH against *TCF3*::*PBX1,* and HeH against *ETV6*::*RUNX1* cases (two-sided Mann-Whitney test). HeH: high hyperdiploid; NS: non significant.

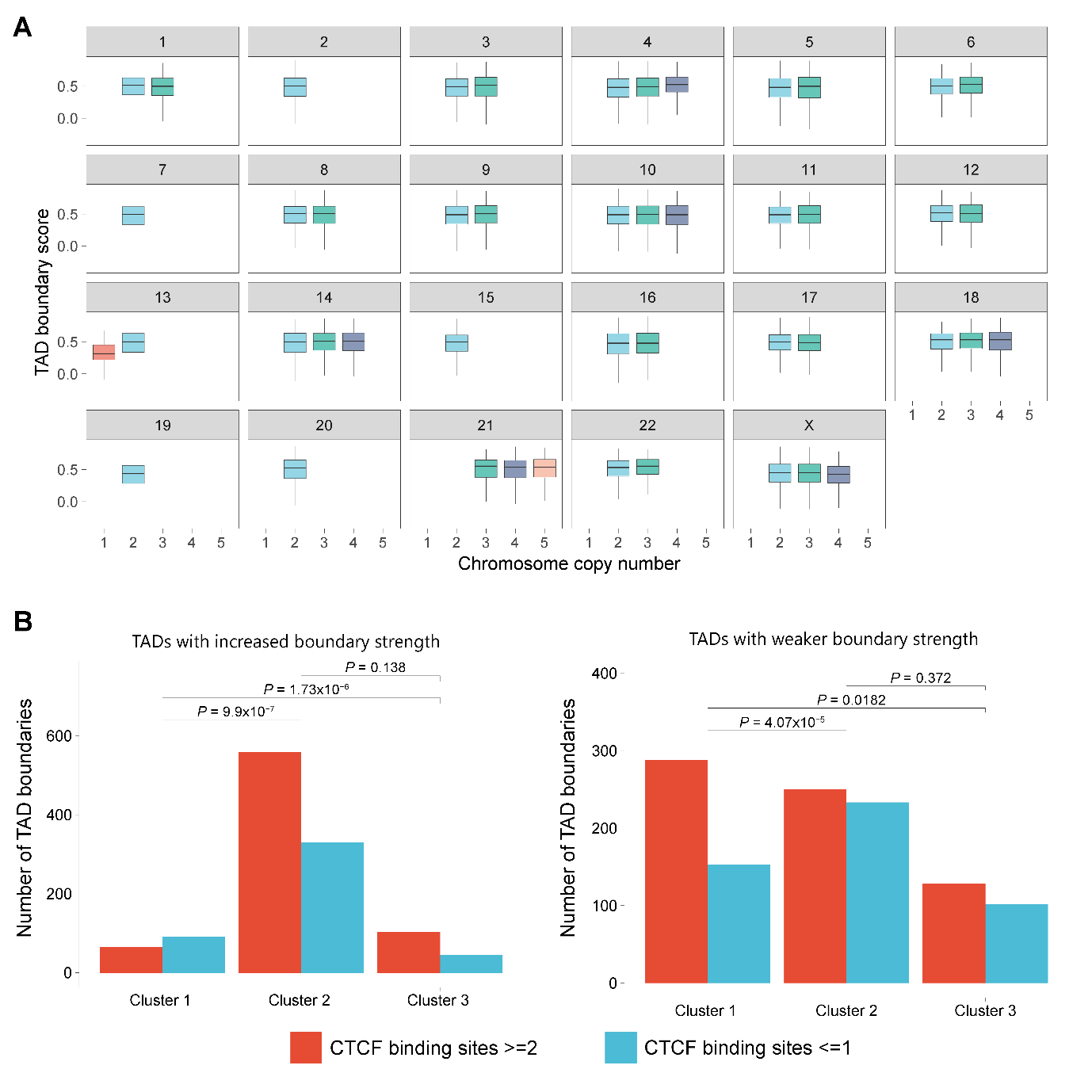

**Supplementary figure 3. Topologically associating domain (TAD) boundaries in different clusters. A.** The median TAD boundary strength for chromosomes 1–22 and X in high hyperdiploid (HeH) cases, stratified by chromosome copy number. No significant differences in TAD boundary strength were observed among chromosomes with varying copy numbers (two-sided Mann-Whitney test). **B.** The left panel shows the relationship between the number of TAD boundaries that exhibited stronger boundary strength in each cluster – when compared with other B-cell precursor (BCP) acute lymphoblastic leukemia (ALL) samples – and the corresponding number of CTCF binding sites. The right panel shows the association between the number of TAD boundaries that displayed weaker strength in each cluster (when compared with other BCP ALL samples) and the number of CTCF binding sites for these boundaries (Fisher's exact test).

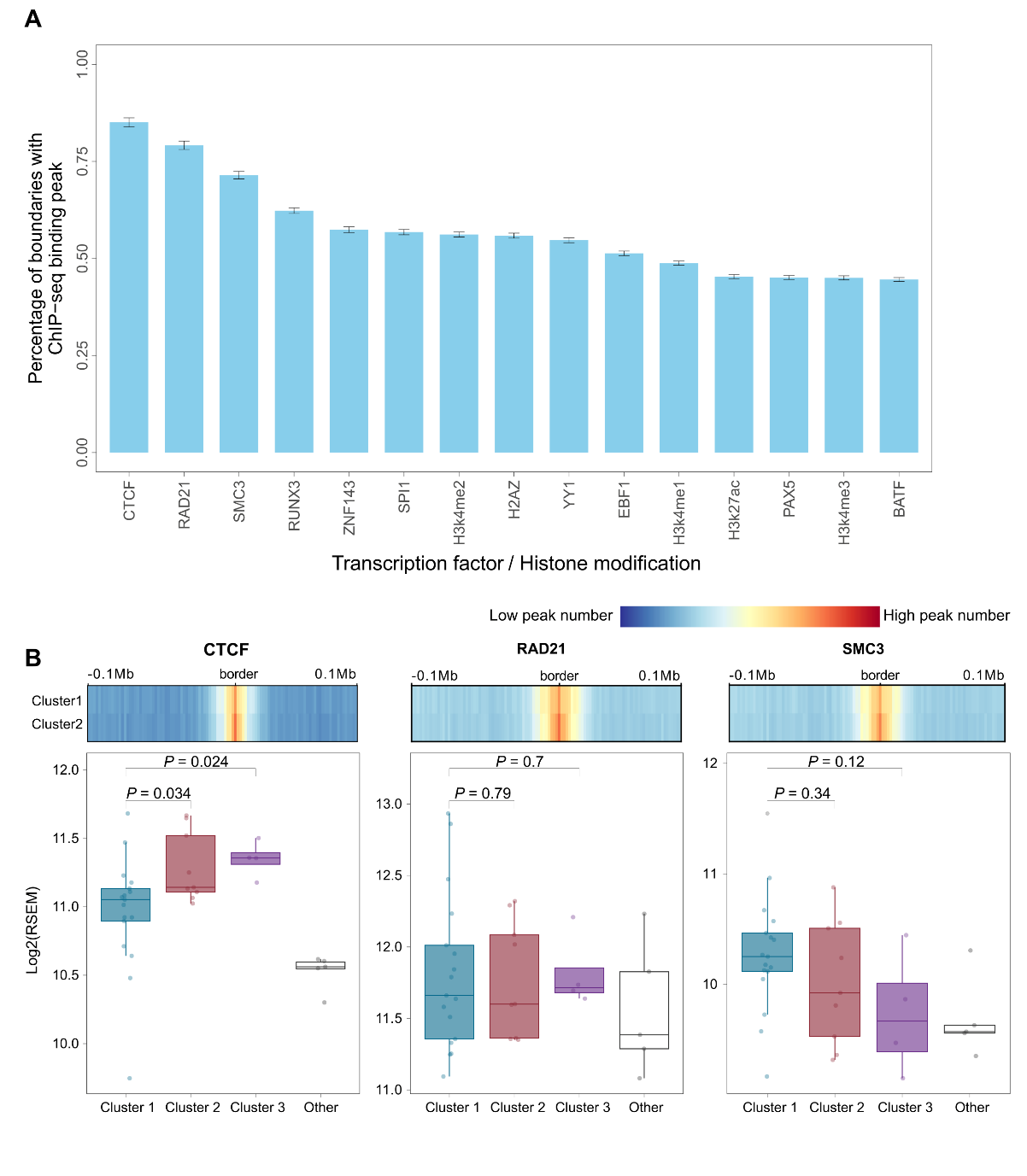

**Supplementary figure 4.** **Transcription factor binding and histone modifications at topologically associating domain (TAD) boundaries.** **A.** The proportion of TAD boundaries that were enriched with key regulatory features, including occupancy by the CTCF/cohesin complex. **B.** Differential analysis of the CTCF/cohesin complex in clusters based on TAD boundaries. In the top sub-panel, the normalized ChIP-seq signals for CTCF, RAD21, and SMC3 are shown at TAD boundaries defined from pooled TAD calls of cluster 1 and cluster 2 cases, respectively. TAD boundaries identified in cluster 2 exhibit higher enrichment of the CTCF/cohesin complex compared with those from cluster 1. The bottom sub-panel displays the expression levels of genes encoding the CTCF/cohesin complex across the different groups. *P*-values were calculated based on two-sided Mann-Whitney test.

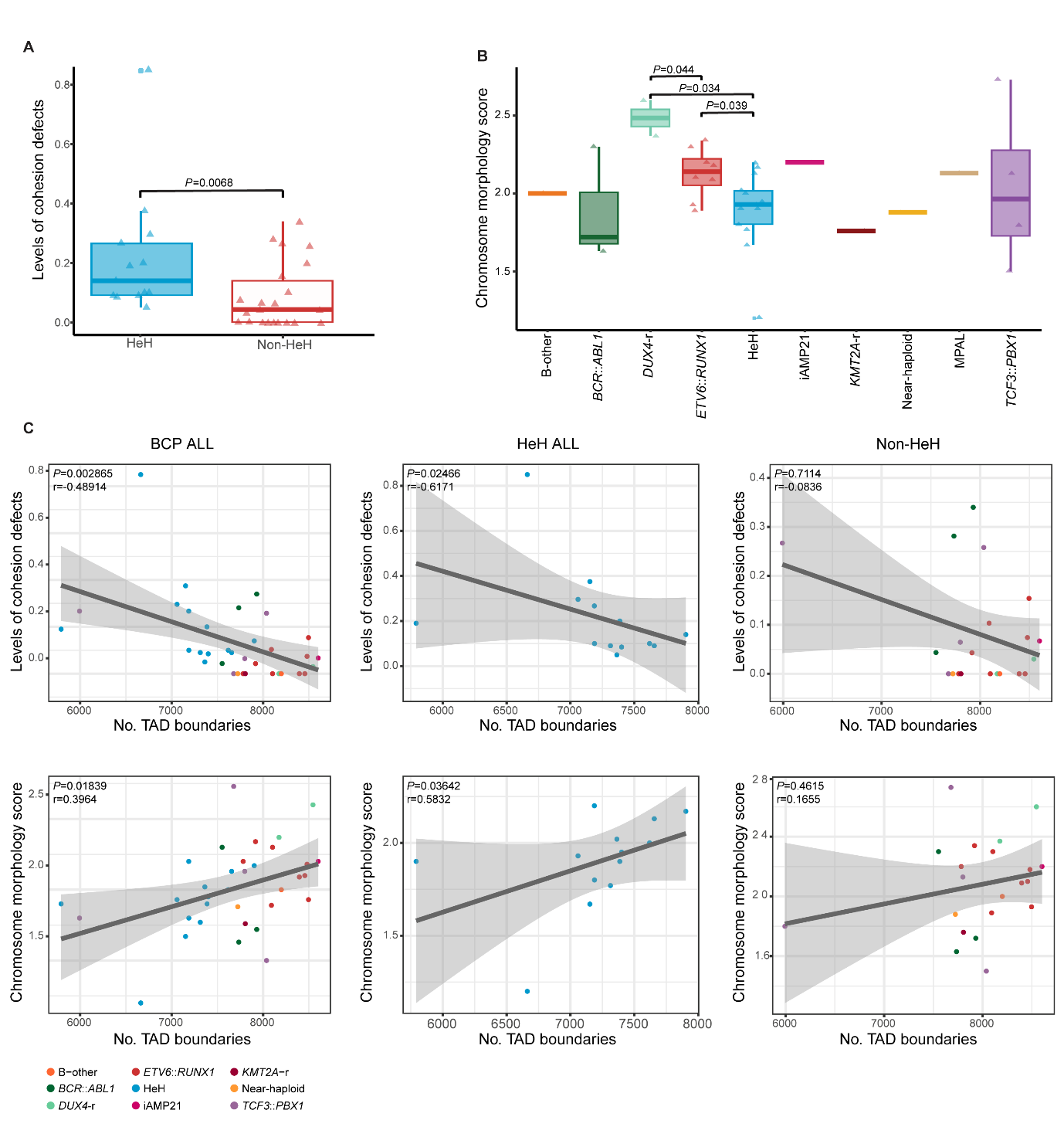

**Supplementary figure 5. Cytogenetic analyses of B-cell precursor (BCP) acute lymphoblastic leukemia (ALL).** Sister chromatid cohesion defects, chromosome morphology score, and comparison between cytogenetic analyses and 3D genomic data. **A.** HeH ALL primary samples display higher levels of cohesion defects compared to other leukemic subtypes (two-sided Mann-Whitney test). **B.** Distribution of chromosome morphology score (scored from 1 – poor to 3 – good morphology) among leukemic subtypes, where the greatest differences were between *DUX4*-rearranged, *ETV6*::*RUNX1* and HeH ALL subtypes (two-sided Mann-Whitney test). **C.** Scatter plots showing the correlation between levels of cohesion defects/chromosome morphology score and number of TAD boundaries (Spearman’s correlation). The analysis was performed considering all BCP ALL cases (left), only HeH cases (center) and excluding HeH cases (right). HeH: high hyperdiploid; iAMP21: intrachromosomal amplification of chromosome 21; MPAL: mixed phenotype acute leukemia; TAD: topologically associating domain.

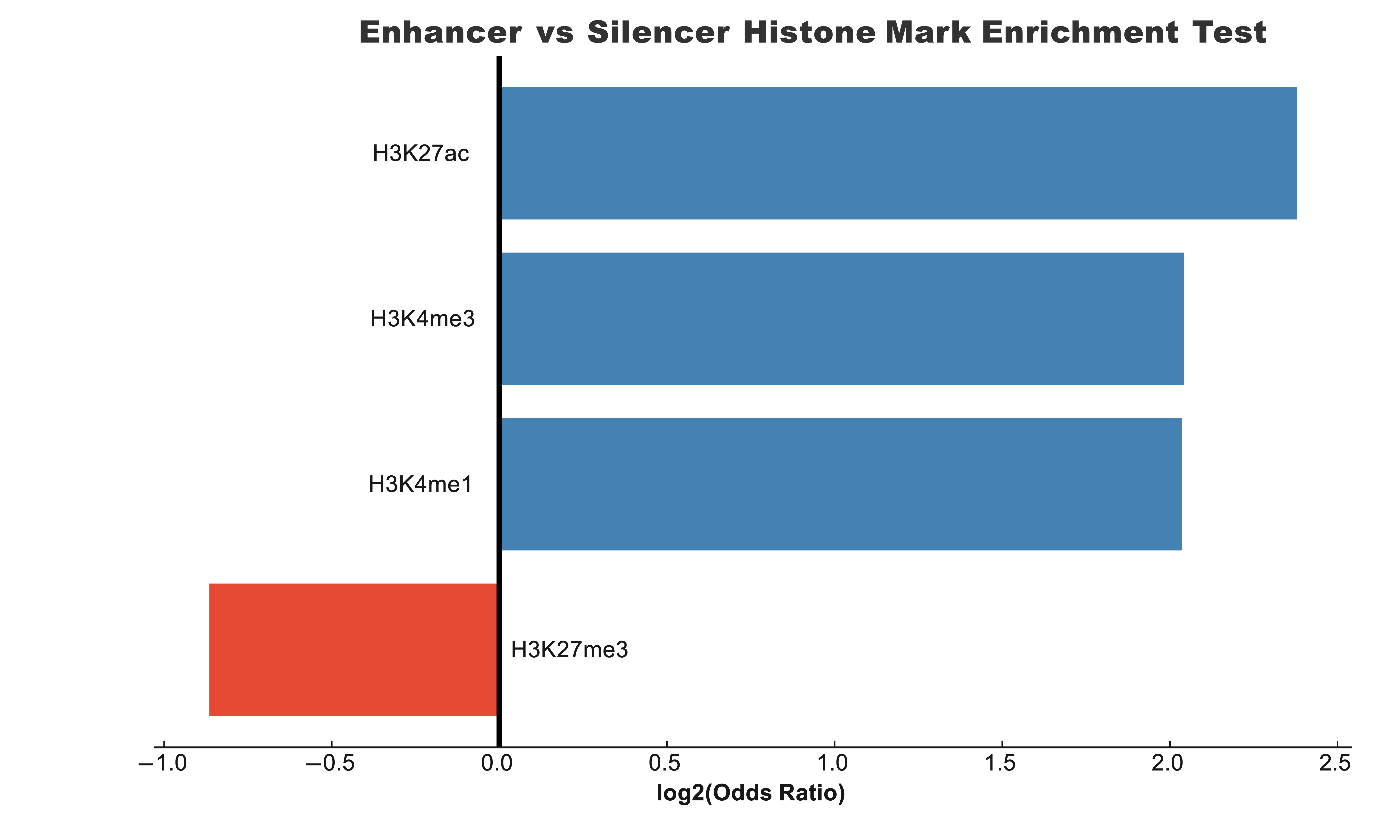

**Supplementary figure 6. Histone mark enrichment results for enhancers versus silencers.** Log2 odds ratios are derived from Fisher’s exact test. Enhancers demonstrate enrichment for H3K4me1, H3K4me3 and H3K27ac while silencers were enriched for H3K27me3.

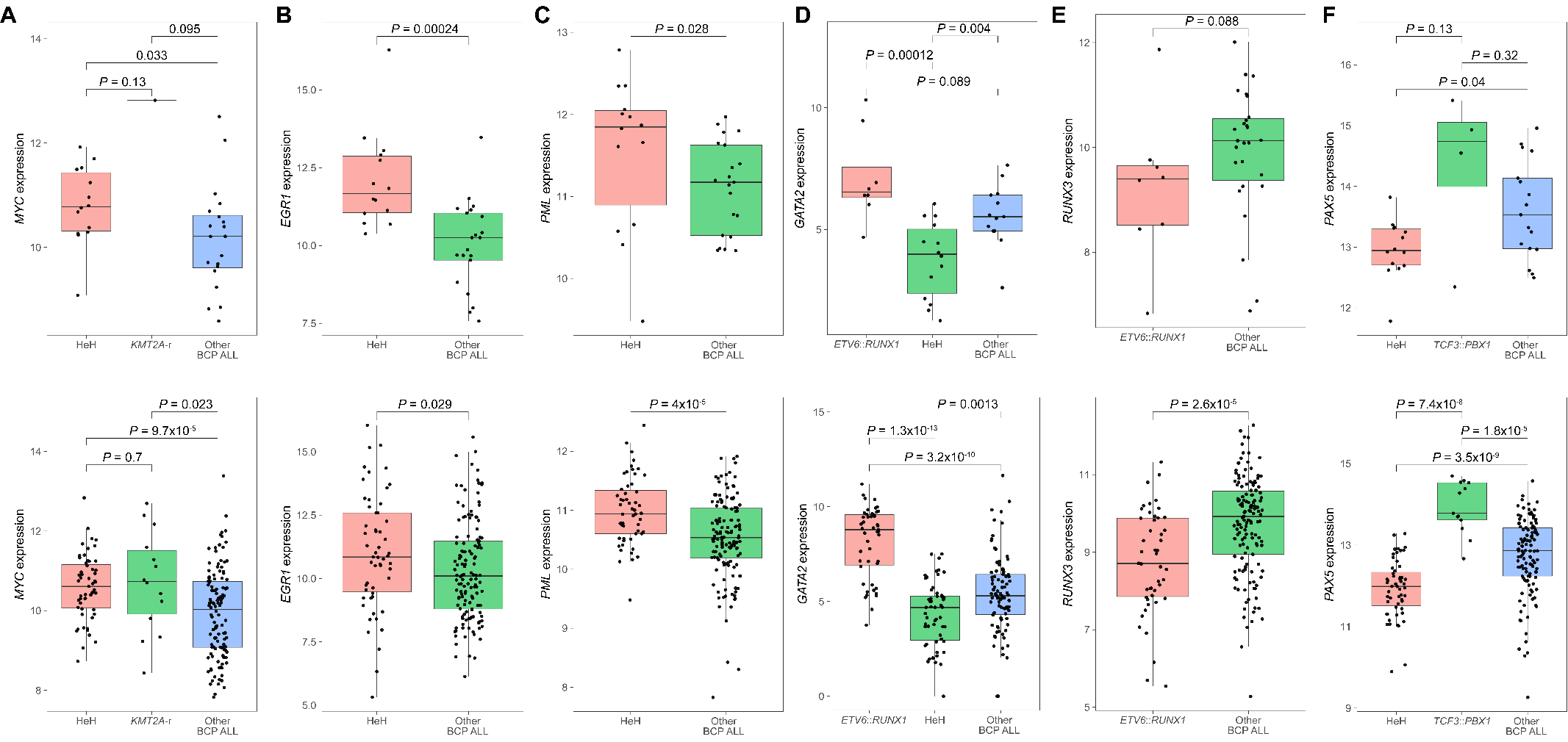

**Supplementary figure 7A-F. Expression of transcription factors in primary B-cell precursor (BCP) acute lymphoblastic leukemia (ALL).** The upper panel shows transcription factor expression levels in BCP ALL samples matched to Micro-C data, and the lower panel shows expression levels in an extended BCP ALL RNA-seq dataset obtained from the European Genome-Phenome Archive (EGA) under accession number EGAS00001001795. *P* values were calculated based on two-sided Mann-Whitney test.

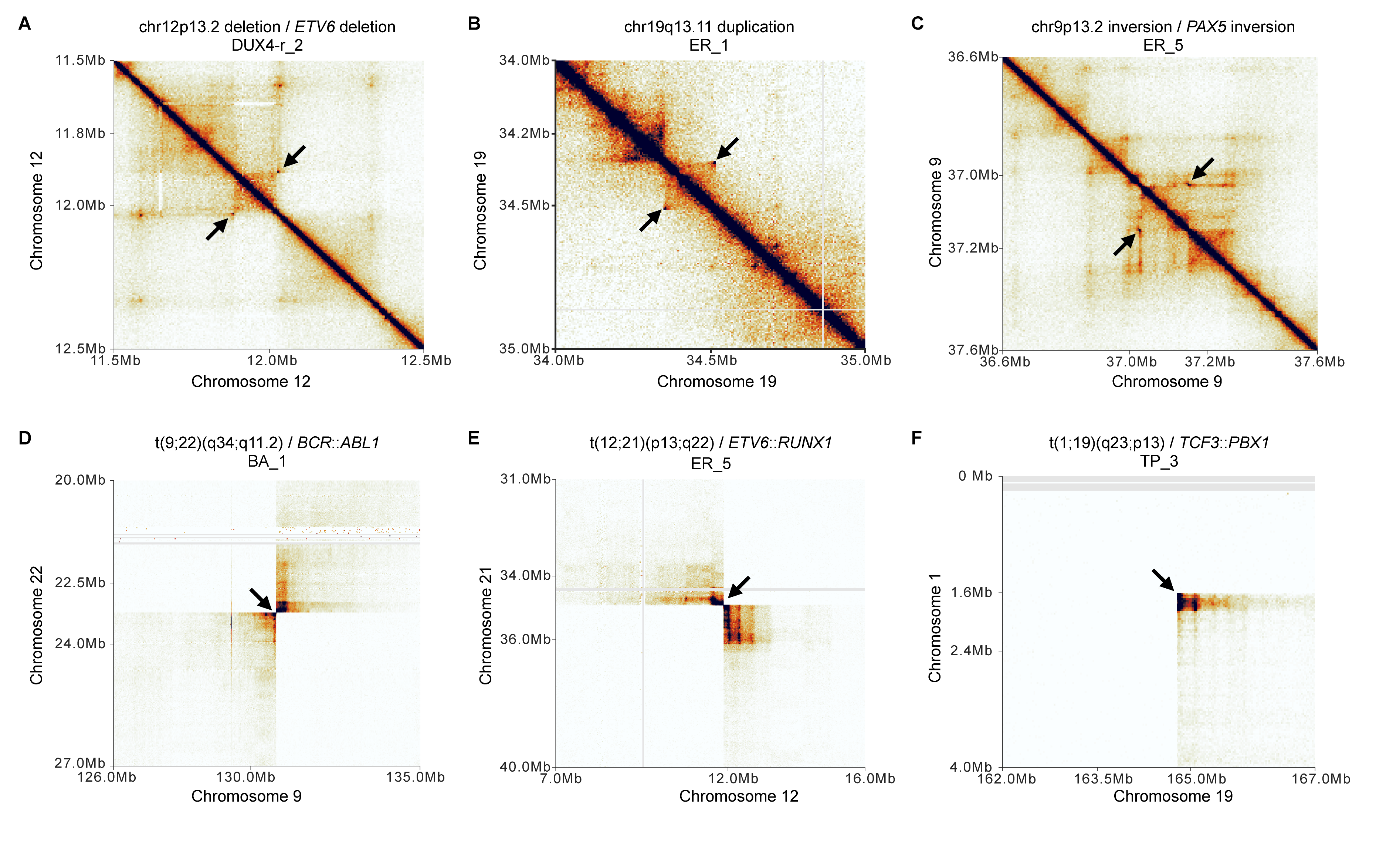

**Supplementary figure 8. Somatic structural variants identified by Micro-C in primary B-cell precursor (BCP) acute lymphoblastic leukemia (ALL).** ICE-normalized Micro-C contact heatmaps showing somatic structural variant subtypes: (A) deletion; (B) duplication; (C) inversion; and (D-F) translocations. Black arrows indicate structural variant breakpoints.

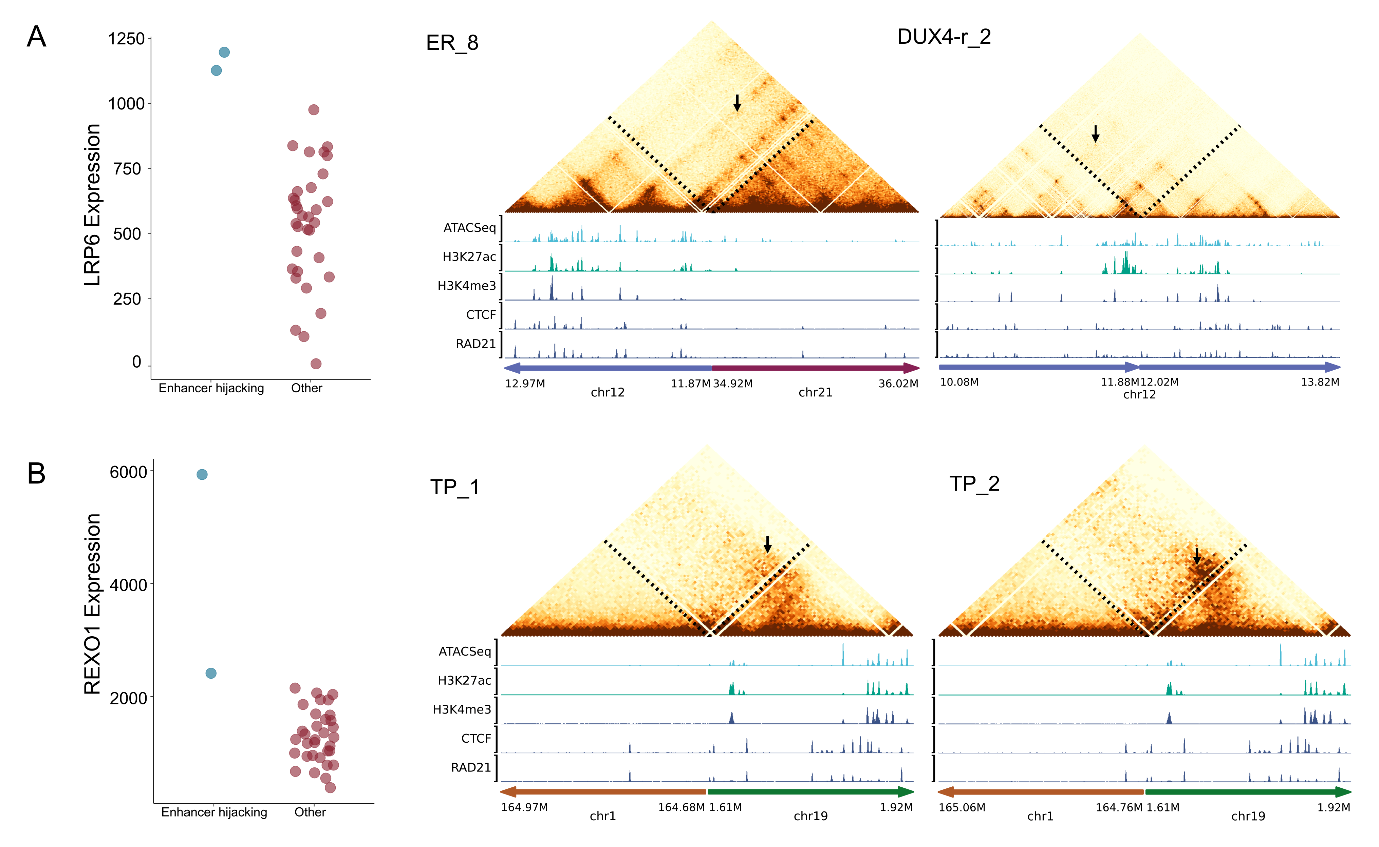

**Supplementary figure 9. Recurrent enhancer-hijacking events driven by somatic structural variants in different B-cell precursor acute lymphoblastic leukemia subtypes.** The left panels show gene expression analysis of enhancer‐hijacking events. Blue dots indicate genes that are involved in enhancer hijacking, whereas red dots represent genes that are not implicated. The right panels show ICE‐normalized Micro‑C contact heatmaps that reveal distinct neoloop formations associated with different somatic structural variants. In the heatmaps, black arrows highlight specific chromatin interactions between gene promoters (marked by blue squares) and the newly hijacked enhancers (marked by yellow squares). The bottom panels present ATAC‐seq and ChIP‐seq tracks for H3K27ac, H3K4me3, and the CTCF/cohesin complex (derived from the GM12878 cell line). **A.** Cases ER_8 and DUX4‑r_2 show neoloops driven by a balanced t(12;21) translocation and a somatic deletion, respectively. **B.** Cases TP_1 and TP_2 display neoloops, detected at 10 kb resolution, that are associated with the t(1;19) translocation.
