## Supplementary tables description for "The 3D genome of pediatric B-cell precursor acute lymphoblastic leukemia"

Description of additional supplementary tables

**ST1.** Patient information including clinical data, supporting data availability and inclusion in previous publications.

**ST2.** Micro-C sequencing metrics and information regarding cluster classification, A/B compartments, topologically associating domains (TADs) and loops.

**ST3.** Shifts in A/B compartments between subtypes, by chromosome, and genes located in the shifted areas. Compartments were quantified as 500 kb bins.

**ST4.** Expression values of 14,560 mRNAs detected in 35 B-cell precursor (BCP) acute lymphoblastic leukemia (ALL).

**ST5.** List of topologically associating domain (TAD) boundary strength of all cases and per cluster.

**ST6.** Summary of cytogenetic analyses, including analysis of sister chromatid cohesion defects and chromosome morphology scoring.

**ST7.** Results from Spearman’s correlation comparing cohesion and condensing genes mRNA expression (Log2(RSEM)) to cohesion defects and chromosome morphology of leukemic samples.

**ST8.** Loops in 1 kb resolution generated from merged Micro-C data along with their corresponding annotations.

**ST9.** Reactome enrichment results for genes with multiple promoter interactions found in Micro-C.

**ST10.** Loop strength score at 10 kb resolution in individual B-cell precursor (BCP) acute lymphoblastic leukemia (ALL).

**ST11.** Detection of structural variants using Micro-C data.

**ST12.** Neoloop and NeoTAD calling result for 35 B-cell precursor (BCP) acute lymphoblastic leukemia (ALL) samples.
